# Sex in the City: Reproductive Behaviors of Dragonflies (Odonata: Anisoptera) across an Urban Landscape

**DOI:** 10.64898/2026.10.13.758078

**Authors:** Christofer Brothers, Aaron Goodman, Ethan R. Tolman, Zander Barnes, Boden Bisgard, Gracie Castillo, Anna Eichert, Kristin Gnojewski, Jim Johnson, Ying Fei Lin, Karie Pappani, Harrison Pardue, Kiichi Tamano, Jessica L. Ware, Christopher D. Beatty

**Affiliations:** University of California, Davis, Department of Department of Neurobiology, Physiology and Behavior, Davis, California, USA; University of Illinois, Urbana-Champaign, Department of Entomology, Urbana, Illinois, USA; American Museum of Natural History, Division of Invertebrate Zoology, New York, New York, USA; Virginia Polytechnic Institute and State University, Department of Biological Science, Blacksburg, VA, USA; Conservation Connection Foundation, Boise, Idaho, USA; Boise State University, Department of Biological Sciences, Boise, Idaho, USA; College of Idaho, Caldwell, Idaho, USA; Vancouver, Washington, USA; Harris Ranch Wildlife Mitigation Association, Boise, Idaho, USA; Graduate Center, City University of New York, New York, New York, USA; Baruch College, Department of Natural Sciences, New York, New York, USA; Southern Illinois University Carbondale, Carbondale, IL USA

**Keywords:** Odonata, Anisoptera, Libellulidae, Aeshnidae, urban ecology, community science, oviposition, tandem mating, hybridization

## Abstract

Anthropogenic cityscapes provide complex habitats that influence animal communication and movement patterns, which are generally understudied in aquatic and semi-aquatic organisms. Community science, referring to public participation in scientific research, is an important tool for recording occurrence records and behavioral observations of understudied urban species. Boise, Idaho (USA), one of the fastest-growing cities in North America, has a long-established community science program for documenting occurrences of the semi-aquatic insect order Odonata (dragonflies and damselflies), making the city an ideal study system for comparing behavioral observations from community science with observations recorded using AI driven iNaturalist. In this study, we test the accuracy of iNaturalist observations and volunteer surveys to characterize dragonfly reproductive behaviors (tandem flight and tandem perching, oviposition, and mating) in the city, and use these observations to determine which dragonflies are using the urban landscape as reproductive habitat. The total numbers of species observations by volunteers and on iNaturalist in Boise were strongly correlated, while reproductive observations showed a strong observational threshold, as reproductive behaviors were rarely identified in less-observed species. iNaturalist tended to underrepresent total occurrences of “flyer” species, which rarely perch, while volunteer surveys underrepresented occurrences and reproductive behaviors of a more cyclical species that did not appear to be abundant in the field season in which this study was conducted. Our findings suggest that Boise provides suitable breeding habitat for both migratory and resident dragonflies. Species distribution models of three migratory and three putatively migratory species further demonstrate that these species would be expected in southwestern Idaho regardless of the presence of a city, and have persisted through urban development. This research demonstrates the value of community science in conducting basic research on understudied urban-dwelling species, although findings must be considered alongside potential biases.

## Introduction

### Urbanization and Odonata Biodiversity in Boise

In the anthropocene, cities provide complex habitats that influence animal communication and movement patterns (Ditchkoff et al., 2006; Delhey & Peters, 2017; Berger-Tal & Saltz, 2019). With city populations increasing rapidly, urbanization poses significant global challenges with profound ecological implications (Halsch et al., 2026). Alteration to land use and the expansion of urban areas have significantly impacted environmental systems (Kalnay & Cai, 2003; Alberti, 2005; Pauleit et al., 2005). Urbanization is one of the main drivers of natural habitat loss, decline in biodiversity, and biotic homogenization (e.g., Piano et al., 2020; Wagner et al., 2021; Halsch et al., 2026). Understanding patterns of biodiversity in the face of urbanization requires baseline data, which unfortunately is lacking for many freshwater macroinvertebrates (Mendonça et al., 2026), such as aquatic insects, for which tolerance to pollution and human disturbance varies greatly across species (Sims, 2014). Further study is needed for identifying which aquatic insects can be found in or near urban waters, and which species have become established in them, using these habitats to complete all, or at least some portion, of their life cycle (Villalobos-Jiménez et al., 2016; Gál et al., 2019).

Community science is a type of study where the public participates in scientific research; community science can expand the spatial and temporal scale of ecological data collection by expanding the number of people doing this work (Bonney et al., 2009), and is a promising means of filling gaps in data about urban insects. Community science can improve our understanding of species distributions and ecological patterns by providing observations from across a greater scale than professional scientists could accomplish on their own (Feldman et al., 2021). Community science can take many forms, from organized volunteers working on targeted research and data collection, to publicly available platforms for data aggregation from contributing observers. One such platform, iNaturalist, has received over 300 million observations from contributors; more than 220 million of these are considered “research grade” (iNaturalist, 2026). The research grade classification generally requires the identification of species to be vetted and confirmed by multiple contributors (Hochmair et al., 2020). However, the biases of these methods and their utility in studying behaviors in urban habitats are poorly understood (Callaghan et al., 2021; Uche-Dike et al., 2026).

### Dragonflies in Boise, Idaho (USA), a model system in urban ecology

Boise, the capital city of Idaho, has over 230,000 residents (U.S. Census Bureau, 2025). The "Treasure Valley" region surrounding Boise has seen record growth in the past few years and is regarded as one of the fastest-growing cities in the U.S. While the population of Boise nearly doubled between 1990 and 2020 (Miller, 2022), little is known about the consequences of this rapid population growth on the surrounding ecosystems and species. In general, increased human activity substantially impacts ecological function and the ecosystem services available in the environment. Boise is traversed by the Boise River, which spans over 6 miles (10 km) through urban, suburban, agricultural, and rural parts of the city. The Boise River Basin has experienced significant anthropogenic changes, including loss of aquatic and riparian habitats (Benda et al., 2003), pollution (e.g., microplastics, Headley et al., 2024; wastewater discharge, Frenzel, 1990; personal care products, livestock fecal steroids, polycyclic aromatic hydrocarbons, and more, Etheridge et al., 2014), logging (MacCoy & Blew, 2005), and higher traffic and recreational volumes than previously recorded as a result of urbanization (Torregrosa & Devoe, 2008; Castro et al., 2018). Despite these changes, urban freshwater habitats in Boise can provide a suitable environment for warm-adapted lentic species (Oertli & Parris, 2019), and, more broadly, urbanization can promote disturbance-tolerant and generalist species (Burkman & Gardiner, 2014). Of the urban aquatic species in Boise, Odonata are amongst the most well studied.

The insect order Odonata, dragonflies and damselflies, is a diverse group with a biphasic life cycle, including an aquatic nymphal stage (Ware, 2021). Different odonate species use different types of freshwater environments; some are well-adapted to life in urban streams and ponds (Tolman et al., 2025), while others are less tolerant of ecological disturbance (Simaika and Samways, 2009). As such, odonates can serve as bioindicators of environmental health (Simaika and Samways, 2009; Simaika and Samways, 2011; Šigutová et al., 2022; Cheri & Finn, 2023). As top predators with broad dietary preferences, Odonata also serve as biocontrol agents of pests in urban ecosystems (Shaalan & Canyon, 2009; Saha et al., 2012). Boise serves as an ideal urban setting for Odonata monitoring with rivers, ponds, wetlands, and reservoirs that support a variety of species (Uche-Dike et al., 2026).

The dragonflies and damselflies of Idaho have been surveyed in a myriad of ways over the last several decades. A particular taxon that resides in Idaho with an expansive range throughout other parts of North America, the blue dasher (*Pachydiplax longipennis* (Burmeister)), was evaluated in Boise, Idaho, as a model species for urban adaptation (Tolman et al., 2025). Few studies have focused on specific species in Idaho since there are no Odonata endemic to Idaho; the majority of studies have instead reported on ecological assemblages. Past community science efforts in Idaho have used “bird count” methodology to document dragonflies and damselflies; a study by 12 observers who visited 3 sites over 3 years in Idaho recorded 27 species of odonates across 6 families (Fothergill & Bouffard, 2006). Comparing environmental DNA (eDNA) sampling of odonates in Idaho to community observations suggests that observers commonly noted species-rich families like Libellulidae but tended to undersample rare taxa, or those flying outside certain time periods when compared to eDNA sampling (Uche-Dike et al., 2026). Despite some observed biases in human observations, Uche-Dike et al. (2026) suggest that community science efforts can capture the majority of species found in the city.

### Community Science and Reproductive Behavior Monitoring

Currently, there are 2.3 million observations of odonates on iNaturalist (iNaturalist, October, 2026). While these are highly useful for distributional data, they are also useful for documenting odonate behaviors. For odonates, presence in a site does not always mean that there is an established, breeding population present, given that these highly mobile insects fly through wide ranges of habitat but may have established breeding populations in only parts of those regions. Observations of dragonflies documenting reproductive behaviors such as “flying in tandem”, “copulating” or “oviposition” can help establish whether species are present, and also whether the environment is a suitable breeding ground with exceptions. This is an important distinction since oviposition often indicates the presence of an established population. However, such reproductive behaviors are not documented in the literature for all species, and when they are, it often comes from a single observation or an unreferenced note in a field guide. Ideally, reproductive behaviors would come from multiple observations taken by several observers, across the period of an adult dragonfly’s lifespan.

### Study objectives

For this study, we relied upon two types of community science data collection methods: iNaturalist observations and volunteer observations. Here, we compare their biases during data aggregation. For example, casual observers of odonates may be more likely to take photographs for iNaturalist of perched versus flying dragonflies, or dragonflies with noticeable color patterning. Using these data we address three questions: *1)* **How useful are iNaturalist/community science photo observations for quantifying and identifying reproductive behaviors in Anisoptera?** *2)* **Which species are using urban Boise as a breeding ground?** And *3)* **Is Boise’s climate more suitable for migratory species than that of the surrounding non-urban landscape?** As we are interested in how the changing landscape affects species distributions and habitat use, we also ask a related question: Does the city provide opportunities for hybridization? We first provide a brief overview of Odonata Ecology, Reproduction, and Migration, which are helpful in contextualizing community science observation of Odonata from Boise.

### Odonata Ecology, Reproduction, and Migration

Odonata comprises ∼6,500 species of freshwater insects (Paulson et al., 2026). Global in their distribution, they are commonly observed at flowing (lotic) and still (lentic) freshwater habitats, whether rural, urban or wild. Dragonflies and damselflies differ in their flight behaviors, with Zygoptera (damselflies) commonly considered perchers, spending the majority of their time perched. Many Anisoptera (dragonflies) are considered to be perchers, but a large number of species are flyers, spending a majority of their time on the wing; some are considered to be intermediate, between perchers and flyers.

Regardless of flight style, dragonflies may be resident or migratory. Migration has been confirmed only in the common green darner (*Anax junius* (Drury)) and the wandering glider (*Pantala flavescens* (Fabricius)), but ∼17 species of North American odonates have been suggested to be putative migrants by the Migratory Dragonfly Partnership (MDP; Migratory Dragonfly Partnership, 2026). In addition to these two species, the MDP lists the spot-winged glider (*Pantala hymenaea* (Say)), black saddlebags (*Tramea lacerata* Hagen), Carolina saddlebags (*Tramea carolina* (Johansson)), striped saddlebags (*Tramea darwini* Kirby; listed by the MDP as *Tramea calverti* Muttkowski), red saddlebags (*Tramea onusta* Hagen), variegated meadowhawk (*Sympetrum corruptum* (Hagen)), autumn meadowhawk (*Sympetrum vicinum* (Hagen)), swamp darner (*Epiaeschna heros* (Fabricius)), taper-tailed darner (*Gomphaeschna antilope* (Hagen)), band-winged dragonlet (*Erythrodiplax umbrata* (Linnaeus)), blue dasher (*Pachydiplax longipennis*), eastern pondhawk (*Erythemis simplicicollis* (Say)), great pondhawk (*E. vesiculosa* (Fabricius)), twelve-spotted skimmer (*Libellula pulchella* Drury) and painted skimmer (*Libellula semifasciata* Burmeister) as putative migrants. Of these, the MDP considers *T. lacerata, P. hymenaea* and *S. corruptum* to be highly likely migrants, and we consider them as migrants here. Some dragonfly species undertake long-distance seasonal migrations that involve different, geographically separated habitats (May, 2013). They can move across extensive geographic ranges, potentially encountering different reproductive environments throughout their migration. Suitable freshwater habitats are important for reproduction. Urban features such as stormwater ponds may provide additional reproductive opportunities (Perron et al., 2024).

Dragonflies and damselflies almost exclusively reproduce in freshwater habitats: females lay eggs and nymphs undergo aquatic development over weeks, months or years, depending on species (e.g., Corbet, 1999). Their reproduction involves characteristic behaviors, including tandem formation, copulation (the mating wheel), and oviposition. Males may or may not guard females, depending on the species; guarding may be either as contact (clasping the female during oviposition, i.e., tandem or interrupted tandem), or non-contact guarding (chasing away males that approach a female; e.g., Corbet, 1999).

Dragonflies and damselflies are commonly observed due to these mating behaviors, but also due to their vibrant color patterns, which may be structural coloration (physical attributes of the cuticle off which light reflects which our eyes perceive as color) or due to pigmentation (e.g., Idec et al., 2024). Wings may be unpigmented, or have yellowish to reddish coloration at the base of the wing. Other species have much more conspicuous wing color patterns, usually with brown or black patches or bands, which are sometimes interspersed with patches of opaque white pigment; some species have entirely, or nearly entirely, opaquely colored wings.

## Methods

### Volunteer training and behavioral observations

We obtained observations from two platforms to evaluate the reproductive behaviors of dragonflies in Boise, Idaho: opportunistic sampling of reproductive observations by harvesting iNaturalist images, and through an organized community scientist program to observe dragonflies in the field. In both platforms, we noted observations of three reproductive behaviors: perching (Fig. 1A) or flying (Fig. 1B) in tandem, copulation (Fig. 1C), and oviposition (Fig. 1D–E). Tandem flight or perching typically occurs before or after copulation (Fig. 1A–B), and in some species it continues while the female oviposits (Fig. 1E; Paulson, 2009). Copulation is when the male, holding the female behind the head with the claspers at the tip of his abdomen, is joined by the female curling her abdomen forward to contact his secondary genitalia on the underside of his second and third abdominal segments, forming the “copulation wheel” (Fig. 1C). Tandem flight is when the male holds the female behind the head with his claspers and the pair flies linked end to end, without genital contact (Fig. 1B). If individuals were observed conducting multiple behaviors simultaneously (e.g. Fig. 1E), both behaviors were noted, although this was only recorded as one observation of a reproductive behavior. As damselfly (Zygoptera) identification often requires extensive expertise, beyond what can reasonably be expected of volunteer community scientists, we restricted observations to dragonflies (Anisoptera).

**Figure 1.**
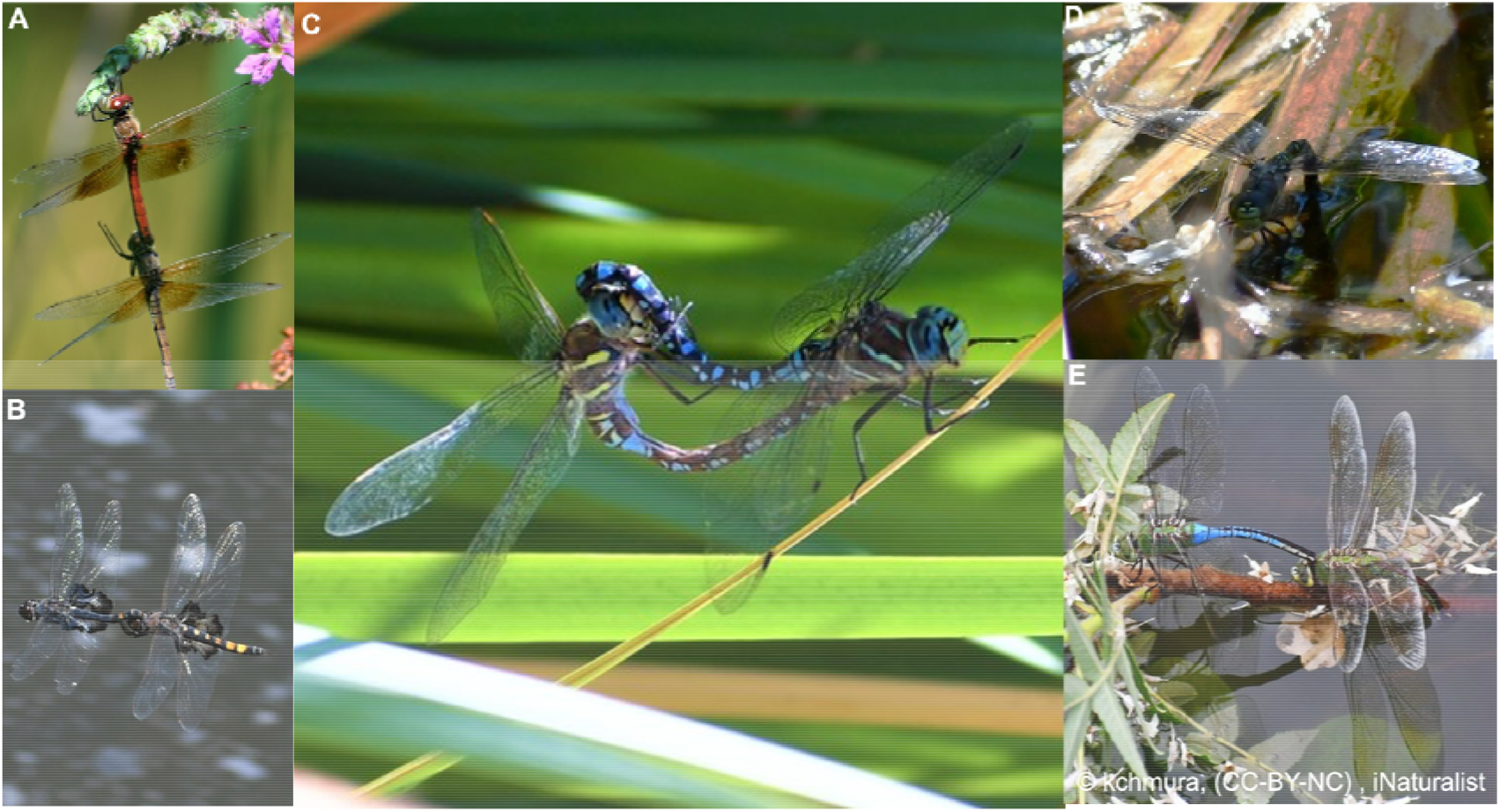
Reproductive behaviors in Anisoptera. Frequencies of tandem pairs, copulation, oviposition, and total reproductive behaviors were modeled separately. Observed dragonflies were coded as being in tandem, in copulation, or ovipositing as follows: (A) Tandem formation during perch, coded as “tandem”; (B) Tandem formation during flight, coded as “tandem”; (C) Copulation/mating wheel, coded as “tandem” and “mating”; (D) Potential solitary oviposition, coded as “oviposition”; (E) Tandem pair at the water surface, potentially contact guarding during oviposition; coded as “tandem” and “oviposition”. For statistical analyses, observations A and B were both recorded as a single observation of a species in tandem, and as one “reproductive behavior” observation per species. C was recorded separately as both a mating and a tandem observation, but was counted as only one “reproductive behavior” observation. D was recorded as an observation of species oviposition, and as one “reproductive behavior” observation. E was recorded separately as both an oviposition and tandem, and included in both models, but was counted as only one reproductive behavior observation. Photographs by Kim Chmura.

### Community scientist monitoring scheme

Nine team members were trained in dragonfly observation, including recognition of species and discrete behaviors. Once trained, volunteers were instructed to make observations at any aquatic site in Boise, and survey for at least 30 minutes. This resulted in surveys of 11 sites between 20 May and 23 September, 2026, where volunteers noted every observed dragonfly and recorded whether each was ovipositing, in tandem, or mating (in copulation).

### iNaturalist observations

We scanned iNaturalist occurrences of Anisoptera in the city of Boise proper as of 1 August 2026, and noted occurrences of species conducting our behaviors of interest. We harvested 1,907 images of Anisoptera by 227 individuals in Boise, Idaho, over the months of June through September, 2010–2026. We reviewed these images and categorized behavior in photographs as ovipositing, in tandem, or mating (in copulation). All species identifications in observations capturing these behaviors were verified by odonatologist Jim Johnson.

Data could not be recorded blindly because our study involved free-living animals observed in the field and in publicly available photographs. Our sample comprised free-flying adult dragonflies recorded opportunistically by iNaturalist users and by trained volunteers at publicly accessible water bodies; the sex, age and condition of individuals were generally unknown. Following the STRANGE framework (Rutz & Webster, 2021), we expect these records to be biased towards conspicuous, perching and easily photographed species, accessible sites and fair-weather conditions, and we model these biases explicitly below.

### Modeling behavioral observations

We analyzed taxa identified to species (n = 28; *Epitheca* sp. was retained as the only record of its genus, and two *Libellula* hybrids were excluded; Table S1). Nomenclature, authorities and common names follow the World Odonata List (Paulson et al., 2026). For each reproductive behavior, and for all reproductive behaviors combined, we compared per-species counts between our two survey platforms, and between each platform and the total number of species observations in Boise on iNaturalist. We also compared total observations between the platforms, and volunteer reproductive observations with total volunteer observations, for a total of 14 count comparisons. In modeling the total number of reproductive behaviors observed per species, each observation was only counted once. Thus, an observation of a species flying in tandem and ovipositing would be included in the models of both of these behaviors, while this observation counted only once in the model of total reproductive behaviors observed per species. We used a generalized linear model (GLM) with a log link and the log(x + 1)-transformed predictor for each comparison. We used a negative binomial (NB) GLM (glm.nb in the R package MASS; Venables & Ripley, 2002) when a likelihood-ratio test indicated overdispersion, and Poisson otherwise. We assessed the strength of association by the deviance explained (D²), Nagelkerke R² (Nagelkerke, 1991) and Spearman ρ, and assessed significance with likelihood-ratio tests against intercept-only models, with Holm correction across the 14 comparisons (Holm, 1979). These count and trait analyses were run in R 4.3.3 (R Core Team, 2024).

We further considered how the behavior and phenotype for each dragonfly species could result in observational bias. For example, species which spend much of their time in flight (flyers) may be less readily photographed than species which spend much of their time perched (perchers). Likewise, a more colorful species may be more readily observed than a more drab one. To capture these potential biases, we modeled each count separately as a function of migratory status (coded by Hedlund et al. [2026] or by the Migratory Dragonfly Partnership [MDP; Migratory Dragonfly Partnership, 2026], in separate candidate models), flight style (percher or flyer, with and without an intermediate class), the presence of any coloration on wings, the presence of wing spots and bands, and family (Libellulidae, Aeshnidae, Gomphidae, and single-species families pooled). Phenotypic trait codings were made according to Lam (2024). Each of the 72 predictor combinations (at most one coding each of migratory status and flight style per model) was fitted as Poisson and NB models and ranked by AICc (Hurvich & Tsai, 1989). The lowest-AICc model was retained; where that model was Poisson but failed its distributional checks, the lowest-AICc NB model was retained instead. We report model support as Akaike weights and relative variable importance (Burnham & Anderson, 2002), and effects as incidence-rate ratios (IRR) with profile 95% CIs. To control for observation frequency, we repeated each behavior model with log(total observations + 1) from the same platform included in every candidate model. We checked model assumptions, including randomized quantile residuals (Dunn & Smyth, 1996), dispersion, simulated zero inflation, Fligner–Killeen homogeneity test, generalized variance inflation factors (car package; Fox & Weisberg, 2019), Cook’s distance, and a family-level random intercept with glmmTMB (Brooks et al., 2017) where family was not already a fixed effect.

### Species distribution modeling of highly observed migratory and putatively migratory species

As we frequently (n > 50) observed six species which are considered to be migratory, or putatively migratory (namely *Anax junius, Sympetrum corruptum*, blue-eyed darner *Rhionaeschna multicolor* (Hagen), *Tramea lacerata*, *Libellula pulchella*, and *Pachydiplax longipennis*; Hedlund et al., 2026; Migratory Dragonfly Partnership, 2026) conducting reproductive behaviors in Boise, we consider whether these species would be expected to be found in southwestern Idaho based on the climate alone, or if they have dispersed to recently suitable urban habitat. For each species, we compared the predicted climatic suitability of the Boise urban area with that of the non-urban landscape within 30 km of the city boundaries (forming an irregular polygon), both as a whole and restricted to areas within 100 m of Boise’s median elevation. Models were fitted separately for each species, following the approach of Tolman et al. (2025) to model habitat suitability in highly observed urban species, using the maximum entropy modeling framework implemented by maxnet (Phillips et al., 2006, 2017) through the ENMeval package in R (Kass et al., 2021; see Supporting Information, Note S1, for details).

## Results

### Wing coloration and patterning predict recorded observations of reproduction

Volunteers completed 55 surveys between 20 May and 23 September 2026, producing 227 taxon records, 48 of which included reproductive behavior. On iNaturalist, we compiled 101 observations of reproductive behavior by 24 observers; the study taxa had 1,907 iNaturalist observations in Boise in total. Volunteers recorded 20 of the 28 species and reproductive behavior in nine, whereas iNaturalist observations documented reproductive behavior in 16 species. Two hybrids between the eight-spotted skimmer (*Libellula forensis* Hagen) and *L. pulchella* were identified on iNaturalist, and one iNaturalist observation showed a tandem pair of two different species, a paddle-tailed darner (*Aeshna palmata* Hagen) and a shadow darner (*Aeshna umbrosa* Walker).

The total number of species observations by volunteers and from iNaturalist were strongly correlated (D² = 0.57, p < 0.001; Fig. S2). The total number of volunteer observations of reproductive behavior was positively correlated with both the number of species observations on iNaturalist (D² = 0.63, p < 0.001) and with the total number of volunteer observations of the species (D² = 0.80, p < 0.001; slope 1.72). Both relationships, however, showed a strong threshold: volunteers rarely recorded reproduction in the less-observed species (Fig. S2). The two platforms agreed moderately in the total number of reproductive observations per species (D² = 0.27, p = 0.025) and tandem behavior (D² = 0.37, p = 0.032). The relationships were not significant for oviposition or mating (p = 0.08 for both). All p-values for these comparisons are Holm-adjusted (Table 1).

**Table 1.** Associations between species-level observation counts (n = 28 species). Each row is a count GLM (log link) with log(x + 1) of the predictor; negative binomial (NB) where a likelihood-ratio test showed overdispersion, otherwise Poisson. Reproductive totals count each record or survey response once per species.

| ID | Response | Predictor | Model | Slope (95% CI) | $\chi^2$ | p | P (Holm) | D<sup>2</sup> | R<sup>2</sup>N | $\rho$ | Flagged species |
| --- | --- | --- | --- | --- | --- | --- | --- | --- | --- | --- | --- |
| *All reproductive behaviors* |  |  |  |  |  |  |  |  |  |  |  |
| C1<sup>†</sup> | Volunteer reproductive | Total iNat | NB | 1.54 (0.84, 2.50) | 18.9 | <0.001 | <0.001 | 0.63 | 0.52 | 0.76 | *S. corruptum*, *T. lacerata* |
| C2 | iNat reproductive | Total iNat | NB | 0.80 (0.41, 1.27) | 12.3 | <0.001 | 0.004 | 0.37 | 0.36 | 0.54 | *A. junius*, *L. forensis* |
| C3 | iNat reproductive | Volunteer reproductive | NB | 0.99 (0.38, 1.70) | 8.2 | 0.004 | 0.025 | 0.27 | 0.26 | 0.50 | *S. corruptum* |
| *Oviposition* |  |  |  |  |  |  |  |  |  |  |  |
| C4<sup>†</sup> | Volunteer oviposition | Total iNat | Poisson | 1.21 (0.74, 1.79) | 37.4 | <0.001 | <0.001 | 0.64 | 0.78 | 0.76 | *A. junius*, *P. longipennis*, *T. lacerata* |
| C5 | iNat oviposition | Total iNat | NB | 1.10 (0.13, 2.44) | 4.5 | 0.034 | 0.082 | 0.25 | 0.17 | 0.30 | *A. junius* |
| C6 | iNat oviposition | Volunteer oviposition | NB | 1.70 (0.09, 3.75) | 3.6 | 0.057 | 0.082 | 0.21 | 0.14 | 0.27 | *A. junius*, *R. californica* |
| *Tandem* |  |  |  |  |  |  |  |  |  |  |  |
| C7<sup>†</sup> | Volunteer tandem | Total iNat | NB | 1.49 (0.57, 3.01) | 10.4 | 0.001 | 0.010 | 0.48 | 0.35 | 0.65 | *T. lacerata* |
| C8 | iNat tandem | Total iNat | NB | 1.22 (0.37, 2.37) | 7.1 | 0.008 | 0.032 | 0.34 | 0.24 | 0.48 | *T. lacerata* |
| C9 | iNat tandem | Volunteer tandem | NB | 1.43 (0.56, 2.59) | 7.4 | 0.006 | 0.032 | 0.37 | 0.25 | 0.64 | *S. corruptum* |
| *Mating* |  |  |  |  |  |  |  |  |  |  |  |
| C10<sup>†</sup> | Volunteer mating | Total iNat | Poisson | 1.32 (0.70, 2.12) | 24.9 | <0.001 | <0.001 | 0.70 | 0.68 | 0.78 | *L. pulchella*, *P. longipennis* |
| C11 | iNat mating | Total iNat | NB | 0.63 (0.26, 1.08) | 8.6 | 0.003 | 0.024 | 0.31 | 0.28 | 0.38 | *A. interrupta*, *S. internum* |
| C12 | iNat mating | Volunteer mating | NB | 1.36 (0.25, 2.64) | 4.9 | 0.027 | 0.082 | 0.19 | 0.17 | 0.21 | *P. longipennis*, *S. corruptum*, *L. forensis* |
| *All observations* |  |  |  |  |  |  |  |  |  |  |  |
| C13 | Total volunteer | Total iNat | NB | 0.86 (0.59, 1.16) | 23.5 | <0.001 | <0.001 | 0.57 | 0.57 | 0.75 | *A. junius*, *P. kurilis*, *S. internum* |
| *Within volunteer data* |  |  |  |  |  |  |  |  |  |  |  |
| C14<sup>†</sup> | Volunteer reproductive | Total volunteer | NB | 1.72 (1.14, 2.45) | 29.6 | <0.001 | <0.001 | 0.80 | 0.69 | 0.81 | *P. longipennis*, *T. lacerata*, *S. semicinctum* |

Slope = coefficient on log(x + 1) (an elasticity). $\chi^2$ and p: likelihood-ratio test against the intercept-only model (1 df); p (Holm): adjusted across all comparisons. D<sup>2</sup>: deviance explained; R<sup>2</sup>N: Nagelkerke R<sup>2</sup>; $\rho$ : Spearman rank correlation. Flagged species: |studentized residual| > 2 or Cook's distance > 4/n. † Curvature test significant (threshold pattern); linear model retained for interpretability.

Family was retained in the best models (Table 2) of the total number of species observations from iNaturalist (χ² = 12.1, p = 0.007) and volunteers (χ² = 18.4, p < 0.001). Species in the pooled single-species families (Cordulegastridae, Corduliidae, and Macromiidae) had 0.04× as many iNaturalist records as Libellulidae (95% CI 0.01–0.21), and volunteers recorded no Gomphidae or species of the pooled families. Family was not retained for any reproductive behavior. The MDP migratory status was retained in the best model for 6 of 8 behavioral counts, while the Hedlund classification was not retained in any model. Migrants had more reproductive records than non-migratory species (e.g., volunteer reproductive IRR 5.5 [putative] and 16 [definite]; Table 3; Fig. 2A). However, after adjusting for observation frequency, migratory status was not retained in any model (Table S4; Fig. 2A). Relative to their observation frequency, flyers were more frequently recorded conducting reproductive behavior than perchers (iNaturalist reproductive IRR 3.5 [1.4–9.5], volunteer reproductive 2.1 [1.1–3.8], volunteer tandem 5.6 [2.5–12.7]), while species with spotted or banded wings had more volunteer oviposition records (IRR 2.7) but fewer iNaturalist oviposition records (0.15). All best models met their assumptions (Tables S2 and S6; Fig. S1).

**Figure 2.**
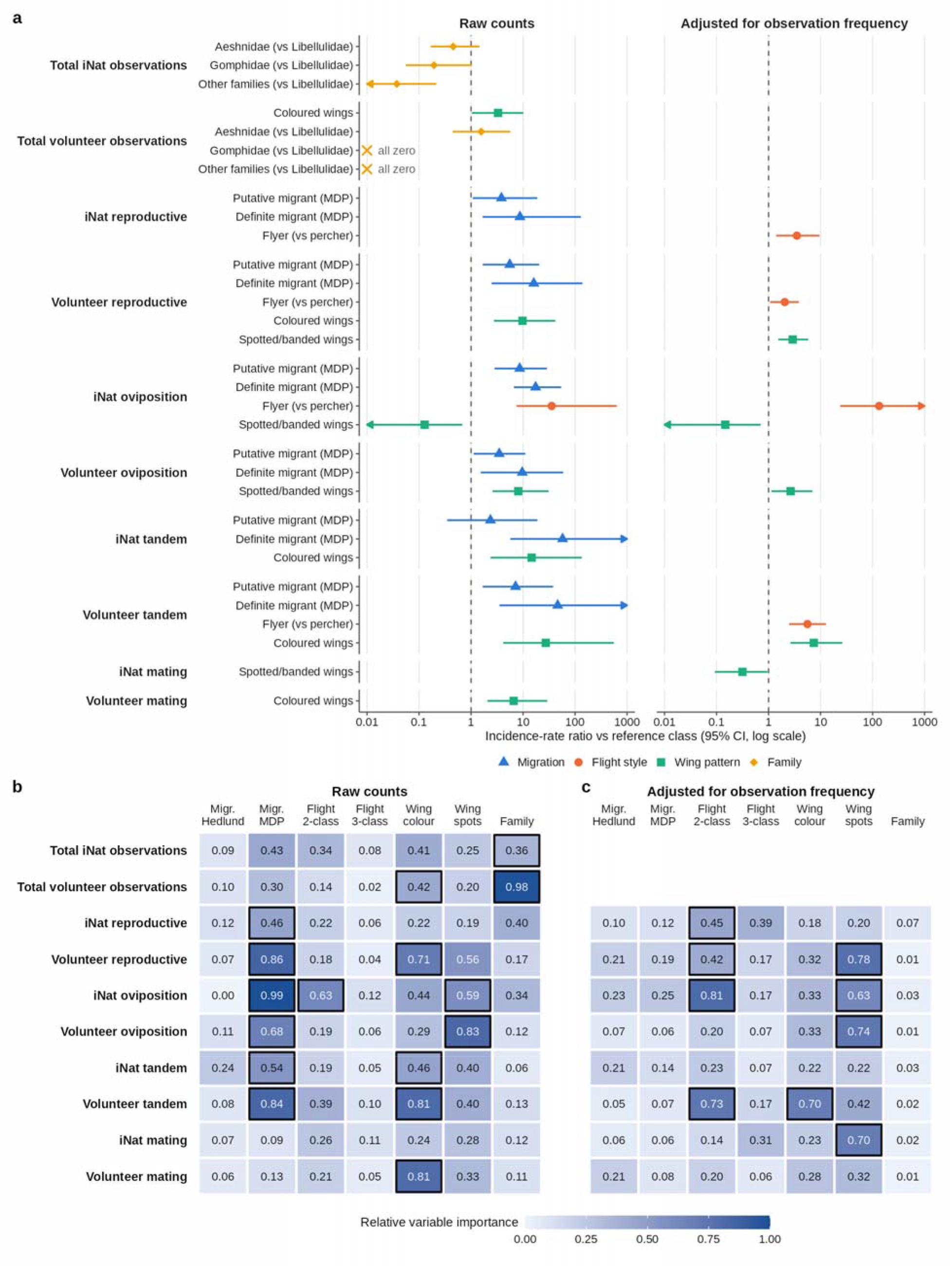
Trait predictors of observation and reproductive-behavior counts across 28 dragonfly species. (A) Incidence-rate ratios (IRR, with 95% profile-likelihood CIs) for the trait terms in the best model for each response. Effects are shown for raw counts (left) and for counts adjusted for observation frequency (right). Each IRR compares a trait class with its reference class: non-migratory, for migratory status from Hedlund et al. or the Migratory Dragonfly Partnership (MDP); percher, for flight style; uncolored wings; unspotted, unbanded wings; Libellulidae, for family. Arrows mark CIs truncated at the axis limits (0.01 and 1000). Crosses mark classes with no observations (“all zero”), whose IRR cannot be estimated; these are plotted at the lower axis limit. Responses whose best model contained no trait terms show no estimate. (B,C) Relative variable importance of each predictor for raw (B) and adjusted (C) counts. Importance is the summed Akaike weight of all candidate models containing that predictor. Outlined cells mark predictors in the retained best model.

**Table 2.**
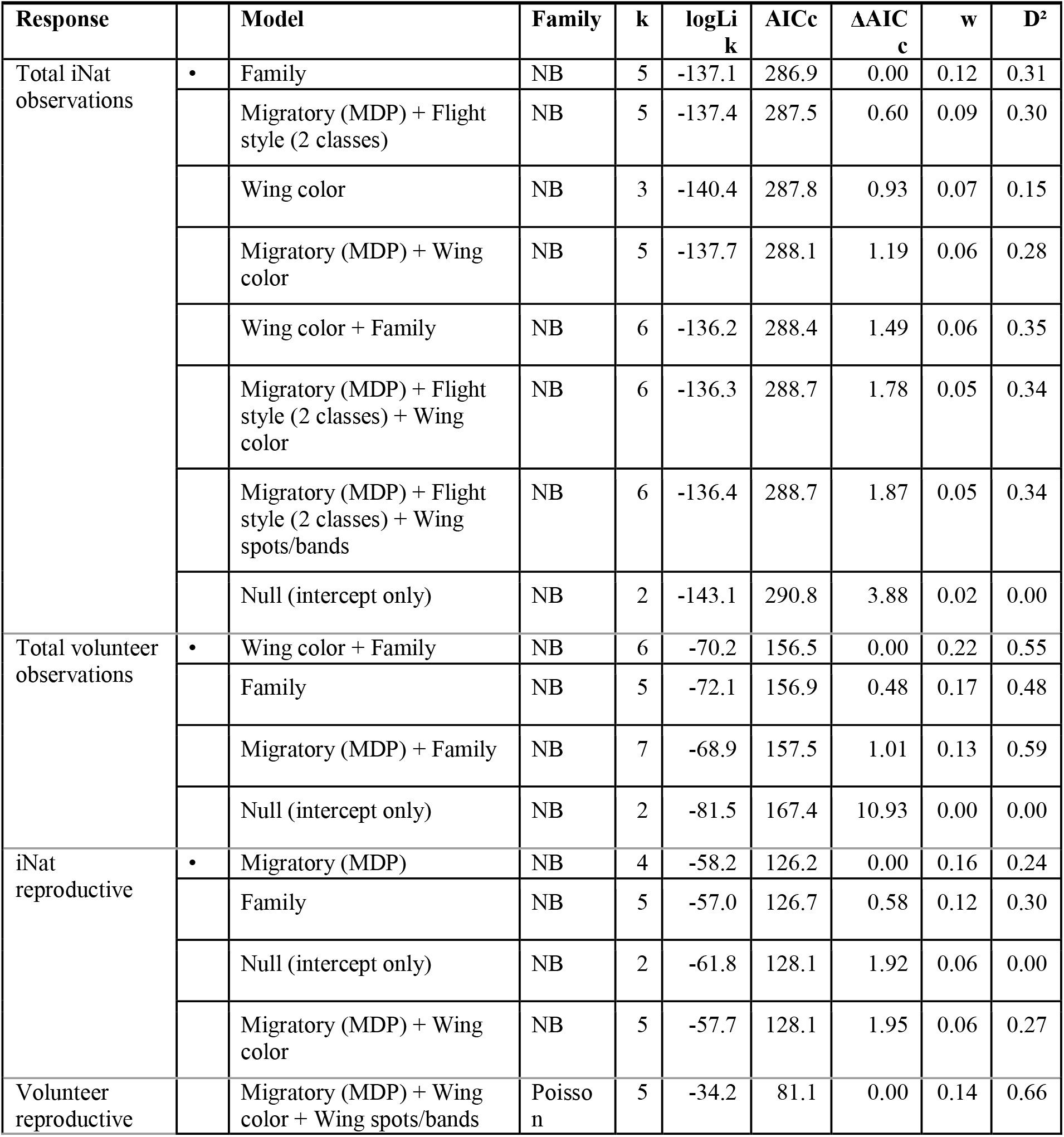
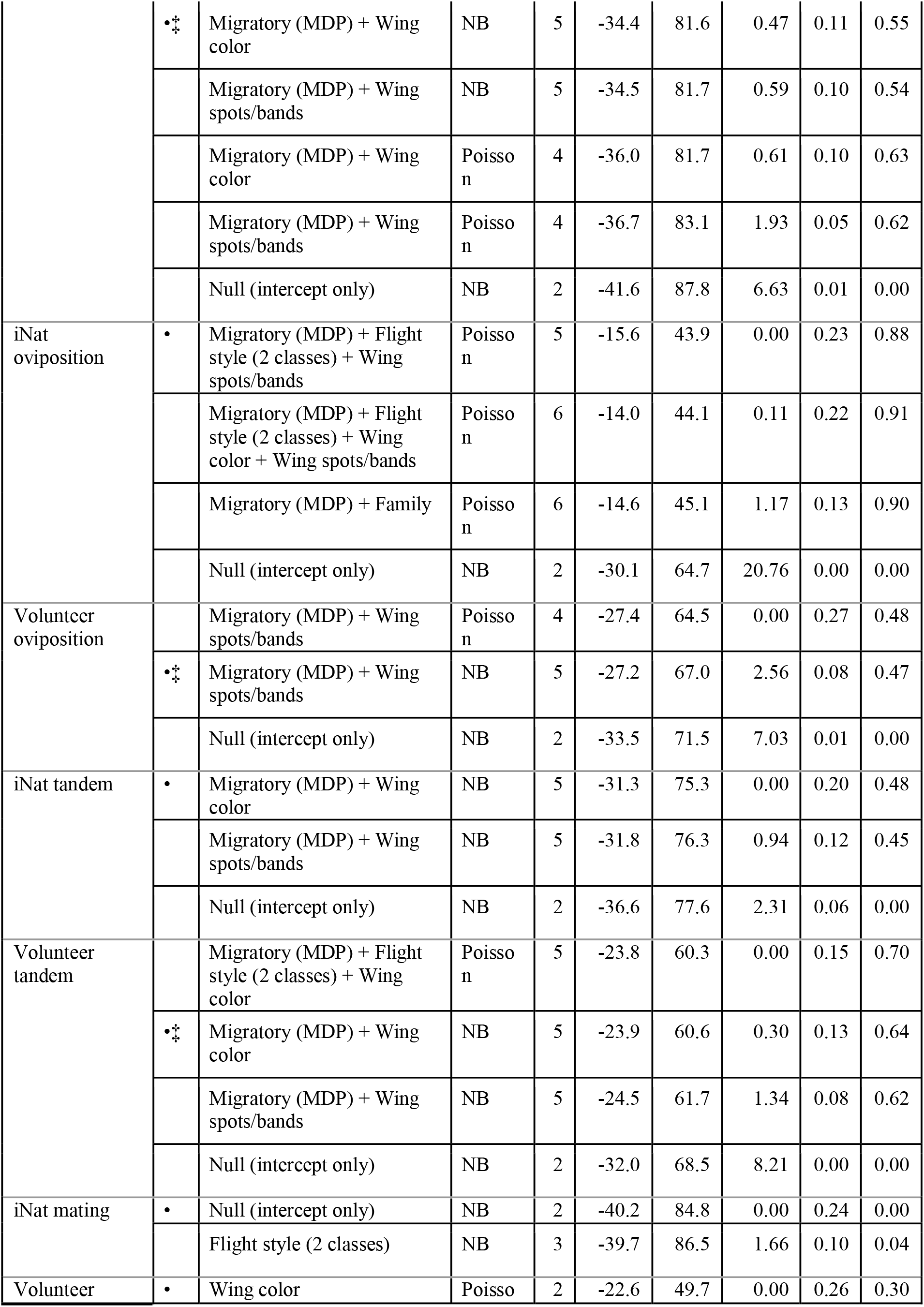
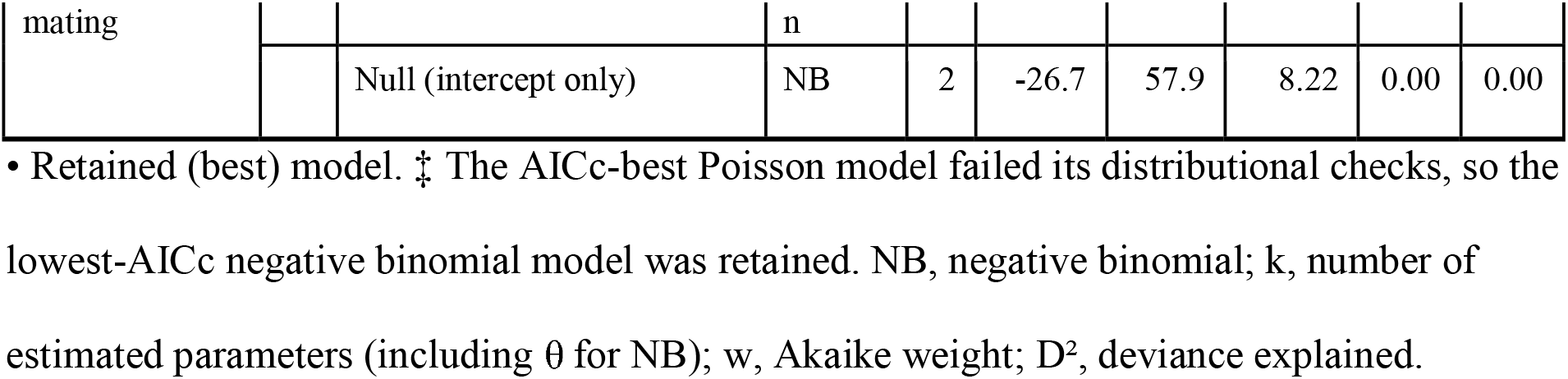
AICc model selection for trait predictors of each observation count (n = 28 species). Shown are all models within ΔAICc ≤ 2 of the best model, and the null (intercept-only) model. Candidate sets combined at most one migration coding, at most one flight-style coding, wing color, wing spots/bands and family, each fitted as Poisson and negative binomial GLMs.

**Table 3.**
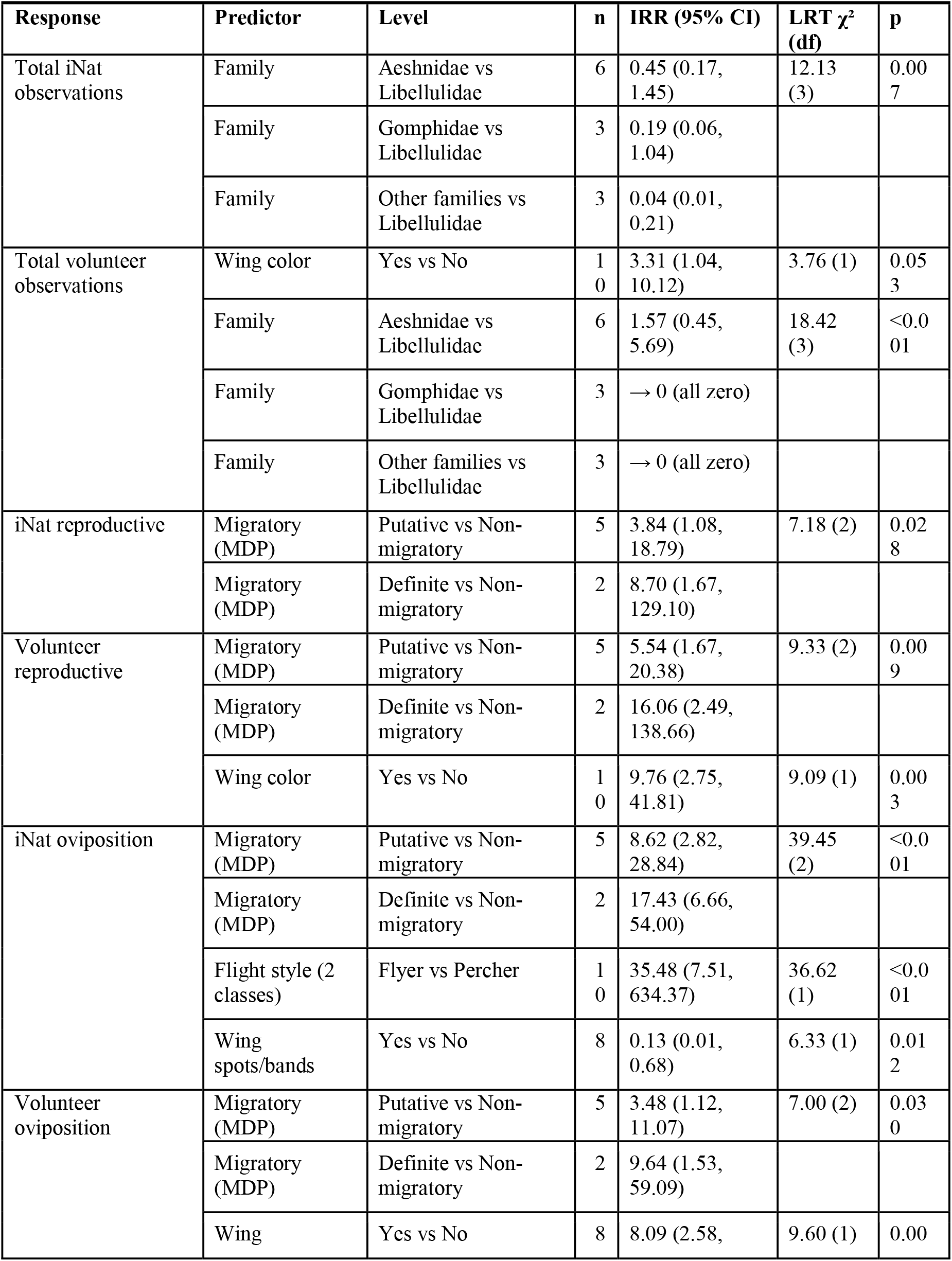
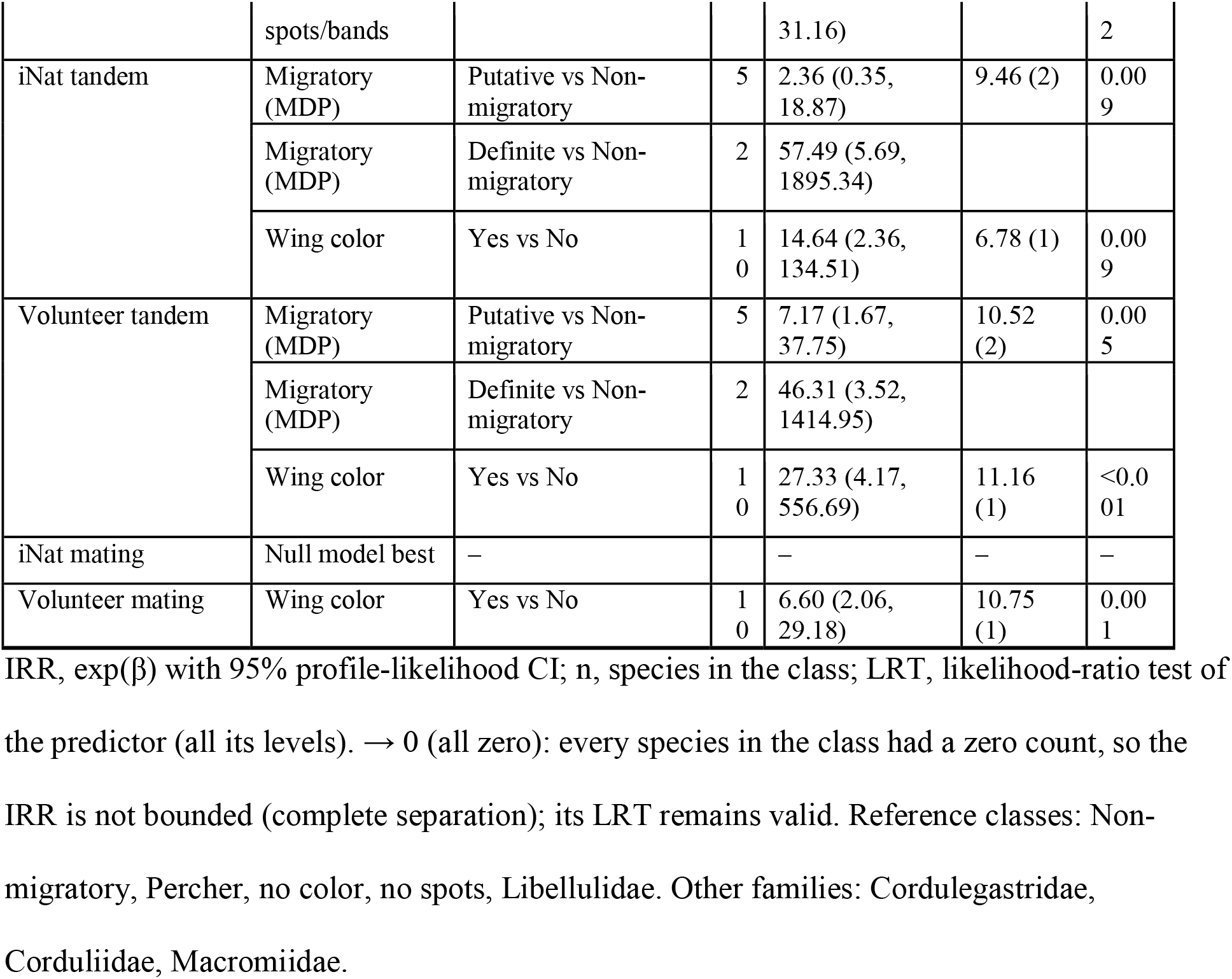
Effects of traits in the best (lowest-AICc) model for each observation count, as incidence-rate ratios (IRR) relative to the reference class.

Several species exhibited higher or lower reproductive behavior than would be expected. The band-winged meadowhawk (*Sympetrum semicinctum* (Say)) had more volunteer reproductive records than expected from its total volunteer observations (C14; Table 1; Fig. S2). *Tramea lacerata* was over-represented in volunteer reproductive and tandem records (Table 1; Fig. S2), and *Anax junius* had more iNaturalist oviposition records than expected (14 vs. 1.7). *Sympetrum corruptum* exhibited the highest disparity between platforms, with 10 iNaturalist reproductive records, and no volunteer reproductive records. *Pachydiplax longipennis* had fewer volunteer reproductive records than expected from its volunteer observations (C14), and fewer volunteer oviposition and mating records than expected from its iNaturalist observations (C4, C10), but more iNaturalist mating records than expected from its volunteer mating records (C12). *Libellula forensis* had no iNaturalist reproductive records despite 197 iNaturalist observations (C2, C12; Table 1; Fig. S2), while the Pacific clubtail (*Phanogomphus kurilis* (Hagen in Selys); Gomphidae) had 48 iNaturalist observations but was never recorded by volunteers. Among the 14 comparisons, removing flagged species changed significance only for iNaturalist oviposition against total iNaturalist observations (unadjusted p = 0.034 to 0.10).

### Species Distribution Modeling

#### Model Fit

After removing exact duplicates, we recovered 23,885 locality records for *Anax junius* (Aeshnidae), 4,714 for *Rhionaeschna multicolor* (Aeshnidae), 18,806 for *Sympetrum corruptum* (Libellulidae), 9,187 for *Tramea lacerata* (Libellulidae), 61,075 for *Pachydiplax longipennis* (Libellulidae), and 11,406 for *Libellula pulchella* (Libellulidae; Table S7). After filtering and thinning, the models were fitted with 2,886 observations for *R. multicolor*, 9,722 for *S. corruptum*, 5,923 for *T. lacerata*, 7,049 for *L. pulchella*, 14,074 for *A. junius*, and 24,804 for *P. longipennis*. The same setting was selected for every species: linear, quadratic and hinge features with a regularization multiplier of 1, which also had the lowest AICc (Akaike weight = 1 for every species). Cross-validated AUC ranged from 0.608 ± 0.008 (*A. junius*) to 0.875 ± 0.009 (*R. multicolor*; Table S7). The models were not overfit (training minus validation AUC ≤ 0.010; omission of held-out records at the 10^th^-percentile threshold was 0.100–0.101) and ranked held-out records well (continuous Boyce index 0.963–1.000; Table S7). All models performed better than null models built from random points (p < 0.001; Table S7).

#### Climatic suitability of Boise and its surroundings

The mean monthly suitability in Boise was at or above each species’ 10^th^-percentile threshold for 3–8 months (*R. multicolor*: Mar–Oct, *A. junius* and *S. corruptum*: Apr–Oct, *L. pulchella*: May–Sep, *T. lacerata*: Jun–Sep, *P. longipennis*: Jun–Aug). Suitability in Boise peaked in June for *L. pulchella* (0.73), July for *S. corruptum* (0.91) and *T. lacerata* (0.73), August for *P. longipennis* (0.61) and *R. multicolor* (0.81), and in September for *A. junius* (0.66). Suitability was low in December–January for all species (0.03–0.16; Table 4; Fig. S7).

**Table 4.**
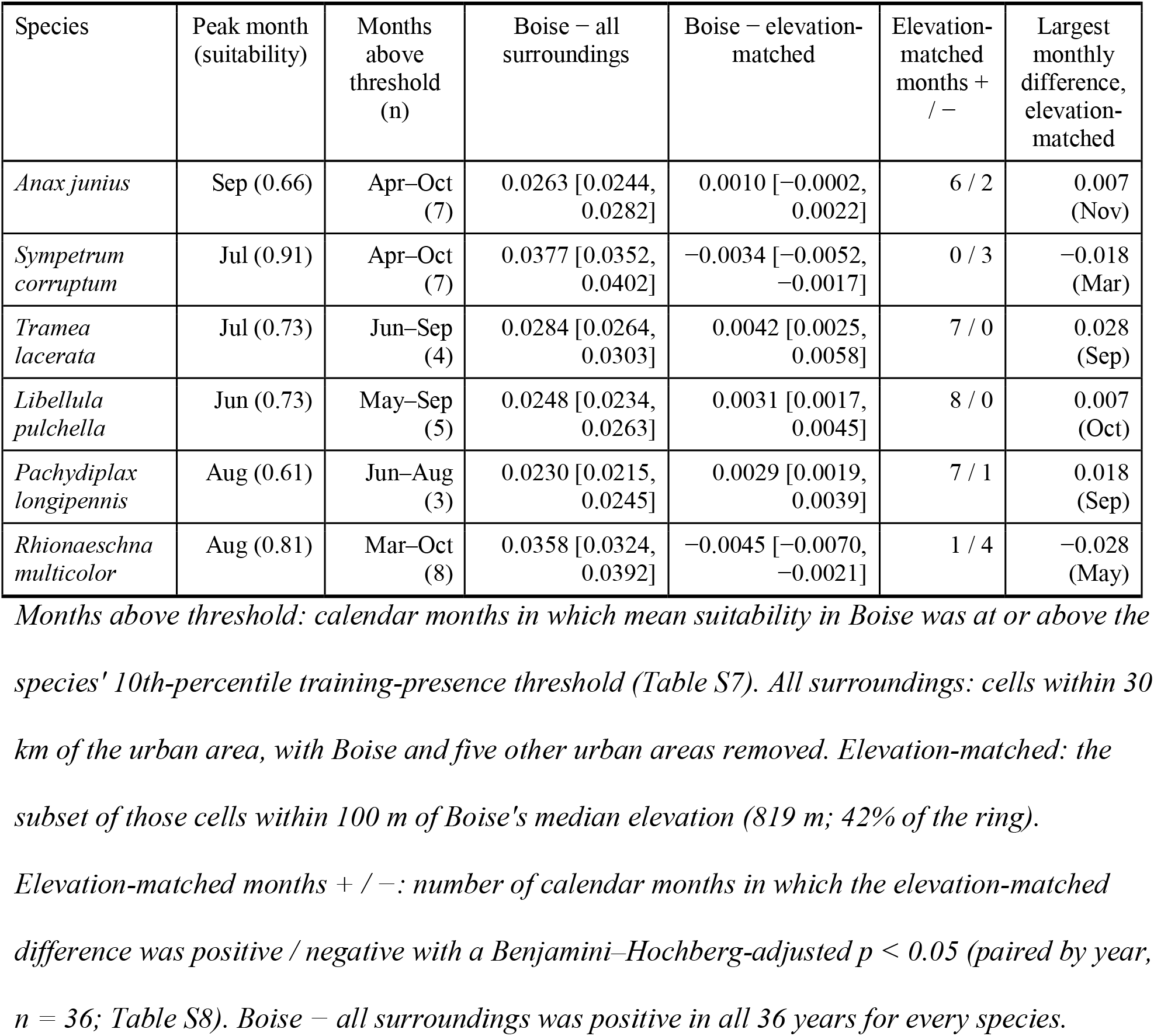
Predicted climatic suitability of the Boise urban area for six dragonfly species, and its difference from the surrounding landscape. Suitability is the long-term (1990–2025) mean of the monthly maxnet predictions (cloglog, 0–1). Differences are Boise minus the comparison area, averaged over the 12 months of each year and then across 36 years (mean [95% CI]).

The area in a 30 km buffer around the boundaries of Boise included 5,338 km² after removing 224 km² of other urban areas, and 2,252 km² of area within 100 meters of the median elevation of Boise (819 m; Fig. S4). Compared with all of its non-urban surroundings, Boise was more suitable for every modeled species in all 36 years when averaged across months (p < 0.01; Table 4), and was more suitable in 10–12 calendar months per species (Benjamini–Hochberg-adjusted p < 0.05; Table S8). The monthly differences were largest in spring (March–June; up to +0.103 for *S. corruptum* in April) and autumn (October–November). In July–August they were near zero, or slightly negative for *A. junius* (July −0.009; Table S8).

The differences between the suitability of Boise and surrounding non-urban elevation-matched regions were about ten times smaller than the difference observed between Boise and all surrounding non-urban areas. Annual mean suitability comparisons ranged from −0.0045 (*R. multicolor*) to +0.0042 (*T. lacerata*), and no calendar month differed by more than 0.028 (Table 4; Fig. 3A). Small positive monthly differences fell mostly in autumn and winter (September–February). For example, Boise was more favorable for *T. lacerata* in September (+0.028) and *P. longipennis* in September (+0.018). Small negative differences fell in spring for *S. corruptum* (March −0.018, May −0.017) and *R. multicolor* (March −0.017, May −0.028; Table S8). Many of these differences in suitability were statistically significant, yet practically minimal at under 3% of the 0-1 suitability scale (Table S8; Fig. 3A).

**Figure 3.**
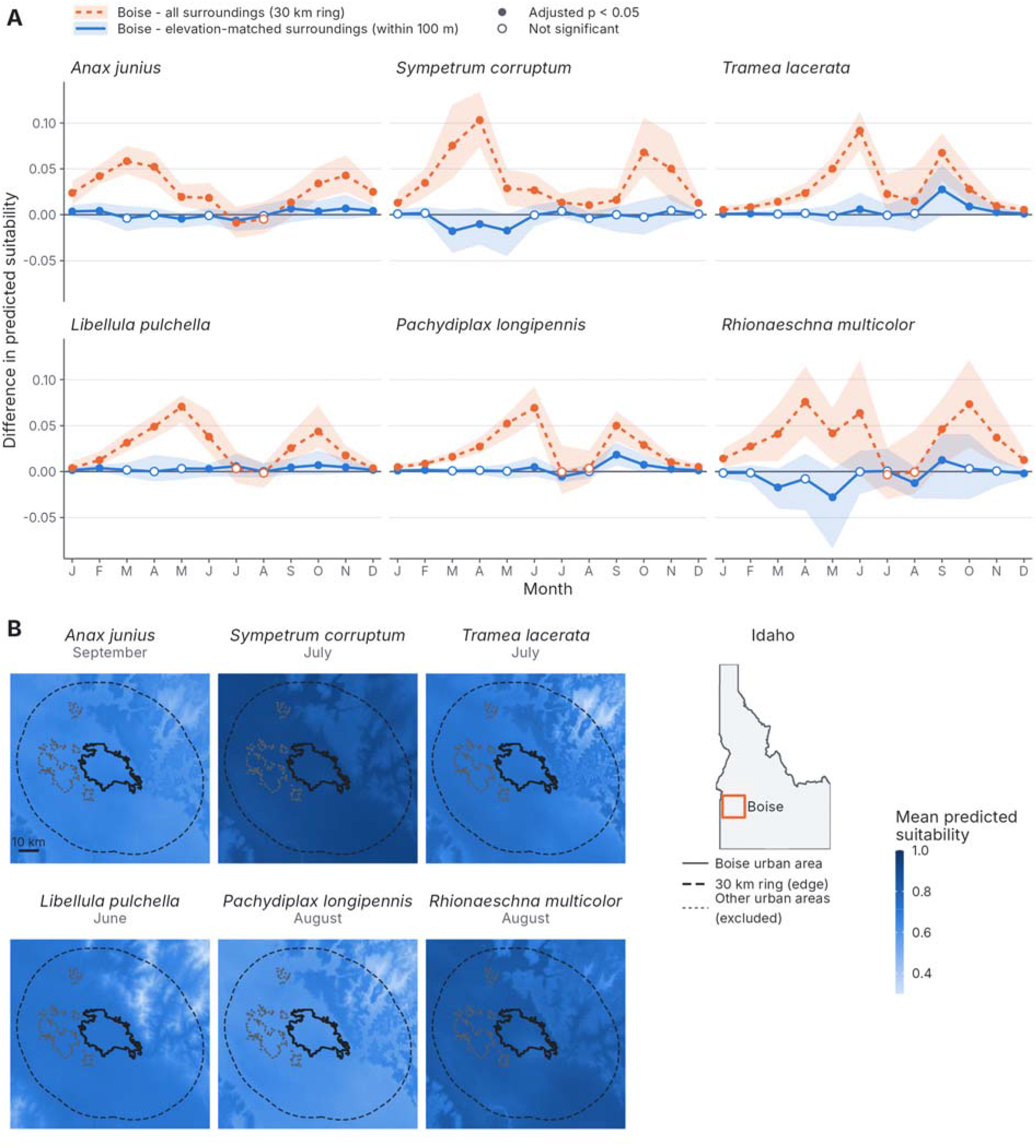
Predicted climatic suitability of Boise, Idaho, compared with its surroundings for six dragonfly species. (A) Difference in predicted suitability between the Boise urban area and its surroundings, by calendar month. Orange dashed lines show Boise minus all surroundings: 800 m cells within 30 km of the urban area, with Boise and five other urban areas (Kuna, Nampa, Emmett, Middleton and Star) removed. Blue solid lines show Boise minus elevation-matched surroundings: the cells of that ring within 100 m of Boise’s median elevation (819 m). Positive values mean Boise is predicted to be more suitable. Lines are means across 36 years (1990–2025), and shaded bands span the 10th–90th percentiles of the yearly differences. Filled points mark months where the yearly differences differ from zero (one-sample t-test, n = 36 years, Benjamini–Hochberg-adjusted p < 0.05); open points are not significant. (B) Long-term mean (1990–2025) predicted suitability around Boise in each species’ peak month in Boise. Solid outline: the Boise urban area (2020 Census “Boise City, ID” urban area, which includes Meridian, Garden City and Eagle). Dashed line: outer edge of the 30 km comparison ring. Dotted outlines: other urban areas excluded from the comparison. Inset: Idaho, with the mapped area boxed in orange. Suitability is the maxnet cloglog output (0–1), from monthly PRISM minimum temperature, maximum temperature and precipitation at 800 m resolution. Values for each area are means of 800 m cells, weighted by the share of each cell inside the area. The color scale starts at 0.3. Scale bar: 10 km.

## Discussion

### Strengths and Biases in Behavioral Observations

The total number of species observations by volunteers and the number of species observations in Boise on iNaturalist were strongly correlated (D² = 0.57, p < 0.001; Fig. S2), while the reproductive observations, as modeled by species observations, showed a strong threshold, as reproductive behaviors were rarely identified in less-observed species (Fig. S2). This threshold could be interpreted in two ways. First, many observations may be needed before reproductive behavior is recorded, because reproduction occupies only a small part of the time odonates spend at the water and some species are rare (Corbet, 1999). Second, some species may reproduce away from the water bodies that observers visit, which would reduce their overall observation counts independently of abundance.

Our two observational methods showed different biases. Here we consider the cases of *Sympetrum corruptum*, *Libellula forensis*, *Pachydiplax longipennis, Tramea lacerata*, and *Anax junius* to explore these biases.

### Sympetrum corruptum

*Sympetrum* that are freshly emerged are drab and often difficult to identify (Paulson, 2009; Lam, 2024), making them difficult to survey via community scientists until they are older and have more noticeable colour patterning. Nonetheless, *S. corruptum* is consistently underrepresented in our volunteer surveys both in general observations and in observations of reproductive behaviors (Table S1; Fig. S2). This highlights a limitation of dedicated volunteer surveys. Such surveys can be difficult to sustain over multiple field seasons, and the dedicated volunteer surveys conducted here spanned only one field season. The City of Boise and partner non-profit institutions first conducted public programs to encourage residents to record dragonfly observations on iNaturalist starting in the spring/summer of 2021 (Uche-Dike et al., 2026). Since this initial push, *S. corruptum* has been recorded on iNaturalist twice in 2021, 10 times in 2022, 65 times in 2023, 22 times in 2024, 19 times in 2025, and 10 times in 2026. Assuming these numbers are somewhat representative of the overall abundance of *S. corruptum* in Boise in a given field season, it is not entirely surprising that *S. corruptum* was not observed by our dedicated volunteers in what appears to be a somewhat down year for the species. Thus, our intensive volunteer survey is not compelling evidence that *S. corruptum* is not using Boise for reproduction, especially as community scientists have recorded a number of reproductive observations on iNaturalist (Table S1; Fig. S2).

### Libellula forensis and Pachydiplax longipennis

While *L. forensis* and *P. longipennis* (both Libellulidae) have both been widely recorded on iNaturalist in Boise, *P. longipennis* was recorded reproducing less often than expected by volunteers, but had more iNaturalist mating records than expected from its volunteer mating records (Table 1; Fig. S2), while *L. forensis* had no records of reproductive behaviors on iNaturalist, but was recorded reproducing by volunteers about as often as expected (Table 1; Fig. S2). As both species are considered perchers (Paulson, 2009), spending much of their time perching on vegetation, they appear to be readily documented on iNaturalist. *Pachydiplax* males have long been noted as territorial (Robey, 1975). Males will defend shoreline territory and actively guard females both in tandem and by aggressively fending off other males (McKinnon & May, 1994) for the span of several hours (Sherman, 1983). As mating behaviors may be occurring in the same localities for several hours, and the intense nature of male-male contests (McKinnon & May, 1994; Baird & May, 2003) may draw human attention, it is plausible that reproductive behaviors in *P. longipennis* are more likely to be noticed by the average iNaturalist user than other species of dragonfly. The reproductive behaviors of *L. forensis* are much less studied, but males are generally not thought to defend a particular territory (Paulson, 2009). The authors have observed that tandem-flights and copulation in this species are very brief, and ovipositing females are not very conspicuous, as they fly low over floating vegetation (although further study is certainly warranted), potentially making their reproductive behaviors less noticeable to the average iNaturalist user.

### Tramea lacerata and Anax junius

*Tramea lacerata* (Libellulidae) and *Anax junius* (Aeshnidae) are both easily identifiable species. *T. lacerata* has distinctive black patterns on the hindwings, while *A. junius* is brightly colored: females and immature males have a bright green thorax, while mature males have a green thorax and a bright blue abdomen (Lam, 2024). As such, they are relatively easy for observers to identify, and they were regularly recorded by our survey team, and a large number of reproductive behaviors were observed.

Despite their conspicuousness, these two species are underrepresented in iNaturalist records. Both of these species are categorized as flyers. Because the flyer mode is characterized by constant flight, often high over water bodies, they are notoriously difficult to photograph. While the necessity of clear images for data contribution and species identification maintains the rigorous objectivity of iNaturalist observations, it can also limit the ability to share observations of more mobile species in formats like iNaturalist, whereas non-photograph documentations such as organized volunteer programs can better document those occurrences in the absence of images (Shirey et al., 2021).

Because both data sources depend on human observers, our samples are likely to be biased towards conspicuous and perching species, accessible urban sites, and fair-weather conditions (Rutz & Webster, 2021). Our conclusions about which species breed in Boise should therefore be read as a lower bound for inconspicuous, high-flying or rarely visited species.

### Urban Boise is reproductive habitat for migratory and resident dragonflies

Although urban spaces have often been regarded as habitats with limited opportunity and resources for breeding odonates (Smith et al., 2022), our findings suggest that Boise provides suitable breeding habitat for both migratory and resident dragonflies. Previous identification of resident *A. junius* populations in urban ponds in the Eastern United States (May & Matthews, 2008) has provided evidence that this species will use urban waters as breeding habitat. Our findings provide further evidence that *A. junius* will use urban habitats for reproduction, while further suggesting that other migratory species (*T. lacerata*, *S. corruptum*) and putative migrants (*P. longipennis*, *L. pulchella*, *R. multicolor*) do the same. Further study observing nymphs and collecting exuvia will be an important next line of evidence in confirming these results.

Our work highlights the potential for cities and urban areas to support tolerant dragonfly populations. Potentially, the increased availability of suitable aquatic habitats (i.e., human-made ponds) in urban regions such as Boise, high dispersal rates among migratory species, and species-specific adaptability to urban landscapes may help describe this pattern. Together, these findings underscore the importance of re-evaluating urban habitats not only as potential barriers to migration, but also as important components of migratory dragonfly ecology and reproduction.

### Hybridization in the city scape

With only one documented occurrence of two different species mating, and two observations of a *L. pulchella* x *L. forensis* hybrid, we do not believe the data presented here provide compelling evidence that the city is serving as grounds for hybridization.

### Boise and surrounding non-urban areas are both suitable habitat for migrants and non-migratory dragonflies

Our species distribution models demonstrated that both Boise and surrounding non-urban regions are broadly suitable (Fig. 3; Table 4). Thus, it does not appear to be the case that our six migrants and putative migrants are only found in Southwest Idaho because urban landscapes provide a haven in an otherwise unsuitable climate (e.g. Southwest Idaho would not be expected to be suitable based on climatic variables, but the species are able to thrive in the city). Rather, they seem to have persisted in the region despite urbanization. We caution that this should not be interpreted as a positive sign for aquatic biodiversity, as high dispersing generalists tend to fare much better than habitat specialists (McKinney, 2006; Clavel et al., 2011; Suhonen et al., 2014). However, it does set up these species as intriguing case studies in urban ecology. What aspects of their biology and physiology allow them to not only be present in an urban area, but use it as a highly productive breeding ground? Such research is urgently needed as we grapple with the effects of human land-use change.

## Conclusion

We find that the most commonly observed dragonflies in Boise, including both migrants and low-dispersing species, are using the city as a mating ground. We further demonstrate the value of community science in conducting such basic research, while cautioning that the use of iNaturalist images and dedicated volunteer surveys are not without biases in conducting behavioral research.

## Acknowledgements

The authors are grateful to all the community scientists in the City of Boise who made this work possible. We are especially thankful for the efforts of Kim Chmura in documenting Odonata, and recruiting and training of community scientists;this work is dedicated to her memory.

## AI Statement

Opus 5.5 (Claude) was used to assist in code generation, and for proofreading the text for minor grammatical errors. All code was manually inspected by the authors, as were suggested grammatical changes.

## Funding

The authors did not receive funding for this study.

## Ethics Statement

This observational study did not involve capture, handling or manipulation of animals, and no permits were required.

## Conflict of Interest

The authors declare no conflicts of interest.

## Data Availability Statement

The data and code that support the findings of this study are openly available in figshare at https://doi.org/10.6084/m9.figshare.34376754. These include the per-species observation and behavior counts, the reviewed iNaturalist records, anonymized volunteer survey records, the filtered GBIF occurrence data, model outputs, and all analysis code. The original GBIF occurrence downloads are available at https://doi.org/10.15468/dl.39422b, https://doi.org/10.15468/dl.4z6pgg, https://doi.org/10.15468/dl.v2b4f3, https://doi.org/10.15468/dl.hyztr9, https://doi.org/10.15468/dl.6gdv97 and https://doi.org/10.15468/dl.45xz9v (GBIF.org, 2026a–f). PRISM climate data are available from the PRISM Climate Group (https://prism.oregonstate.edu), and iNaturalist observations are publicly available at https://www.inaturalist.org.

## Author Contributions

Christofer Brothers, Aaron Goodman, Ethan R. Tolman and Christopher D. Beatty: Conceptualization, Methodology, Investigation, Writing – original draft, Writing – review & editing. Karie Pappani and Kiichi Tamano: Investigation, Writing – original draft, Writing – review & editing. Zander Barnes, Boden Bisgard and Kristin Gnojewski: Investigation. Gracie Castillo, Anna Eichert, Jim Johnson, Ying Fei Lin, Harrison Pardue and Jessica L. Ware: Writing – original draft, Writing – review & editing.

## Supporting Information

Note S1. Species distribution modeling methods.

Table S1. Species included in the analyses, with trait classifications and observation counts.

Table S2. Assumption checks for the best trait model of each observation count.

Table S3. AICc model selection for behavior counts adjusted for observation frequency.

Table S4. Trait effects in the best observation-frequency-adjusted models.

Table S5. Sensitivity of the best raw-count models to the coding of species not listed by the Migratory Dragonfly Partnership.

Table S6. Fit and assumption checks for all 32 count models.

Table S7. Occurrence data, selected settings and evaluation statistics for the six species distribution models.

Table S8. Long-term mean predicted suitability by calendar month in Boise and its surroundings, with paired differences.

Table S9. Mean predicted suitability for every species, year and month (Excel file).

Figure S1. Fit and assumption checks for all 32 count models.

Figure S2. Associations between species-level observation counts for the 14 comparisons.

Figure S3. Observation counts per species by migratory status (MDP).

Figure S4. Comparison areas and elevation matching around Boise.

Figure S5. Response curves of the selected species distribution models.

Figure S6. Long-term mean predicted suitability across Idaho in each species’ peak month in Boise.

Figure S7. Long-term mean predicted suitability around Boise in each calendar month, by species (S7a–f).

Figure S8. Sensitivity of the Boise comparison to the background design. Data S1. Data and R code for all analyses (Supplemental_data).

## Notes

### Competing Interest Statement

The authors have declared no competing interest.

