## Supplemental Figures for "Sex in the City: Reproductive Behaviors of Dragonflies (Odonata: Anisoptera) across an Urban Landscape"

**Supporting Information Figures**

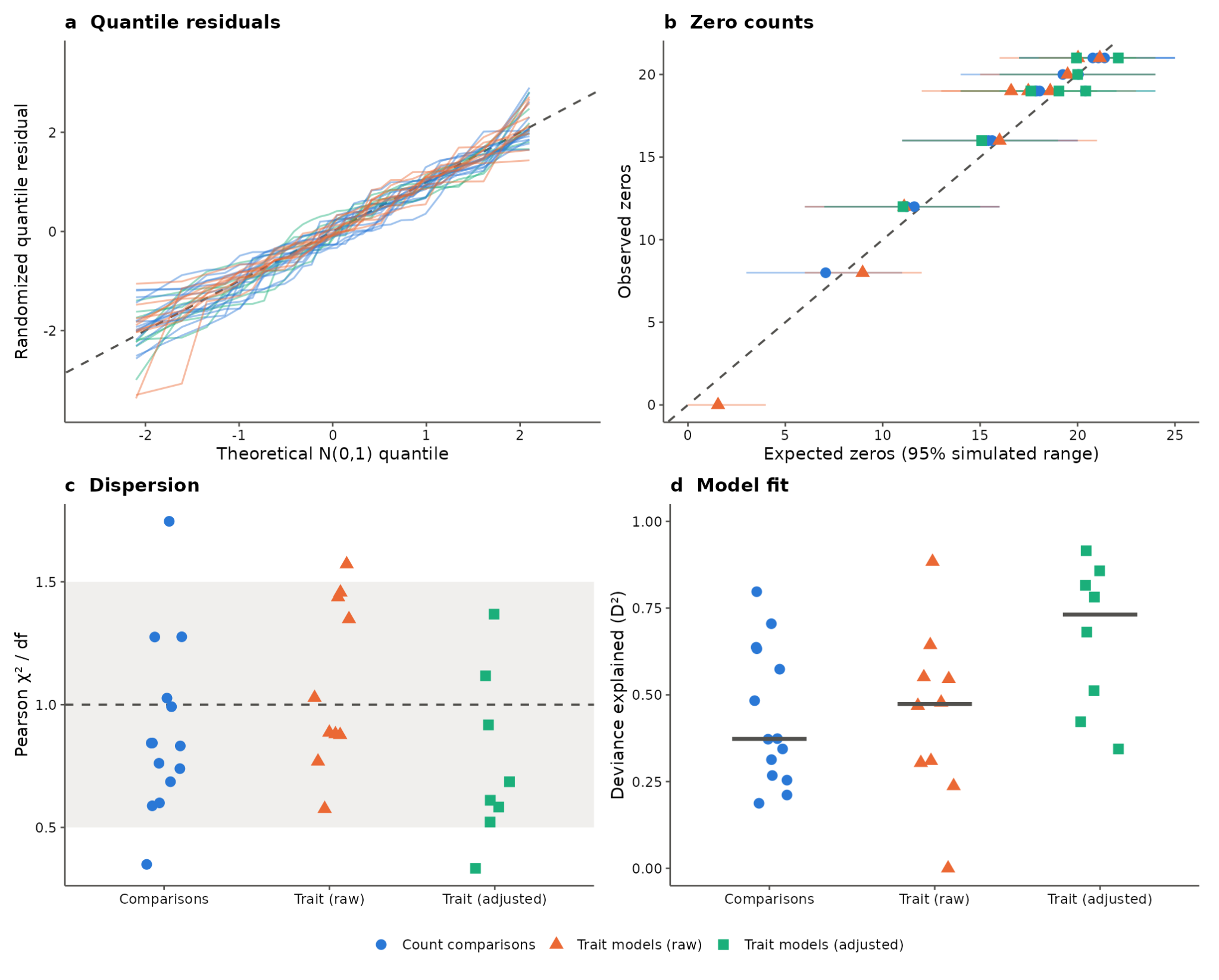

**Figure S1.** Fit and assumption checks for all 32 count models: count comparisons (circles), raw-count trait models (triangles) and observation-frequency-adjusted trait models (squares). (A) Randomized quantile residuals against N(0,1) quantiles, one line per model; the dashed 1:1 line indicates an adequate error distribution. (B) Observed vs. expected number of zero counts, with the 95% range from 2,000 simulations of each fitted model; the dashed line is 1:1. (C) Dispersion ratio (Pearson χ²/df); the shaded band marks 0.5–1.5 and the dashed line marks 1. (D) Deviance explained (D²); bars show medians.

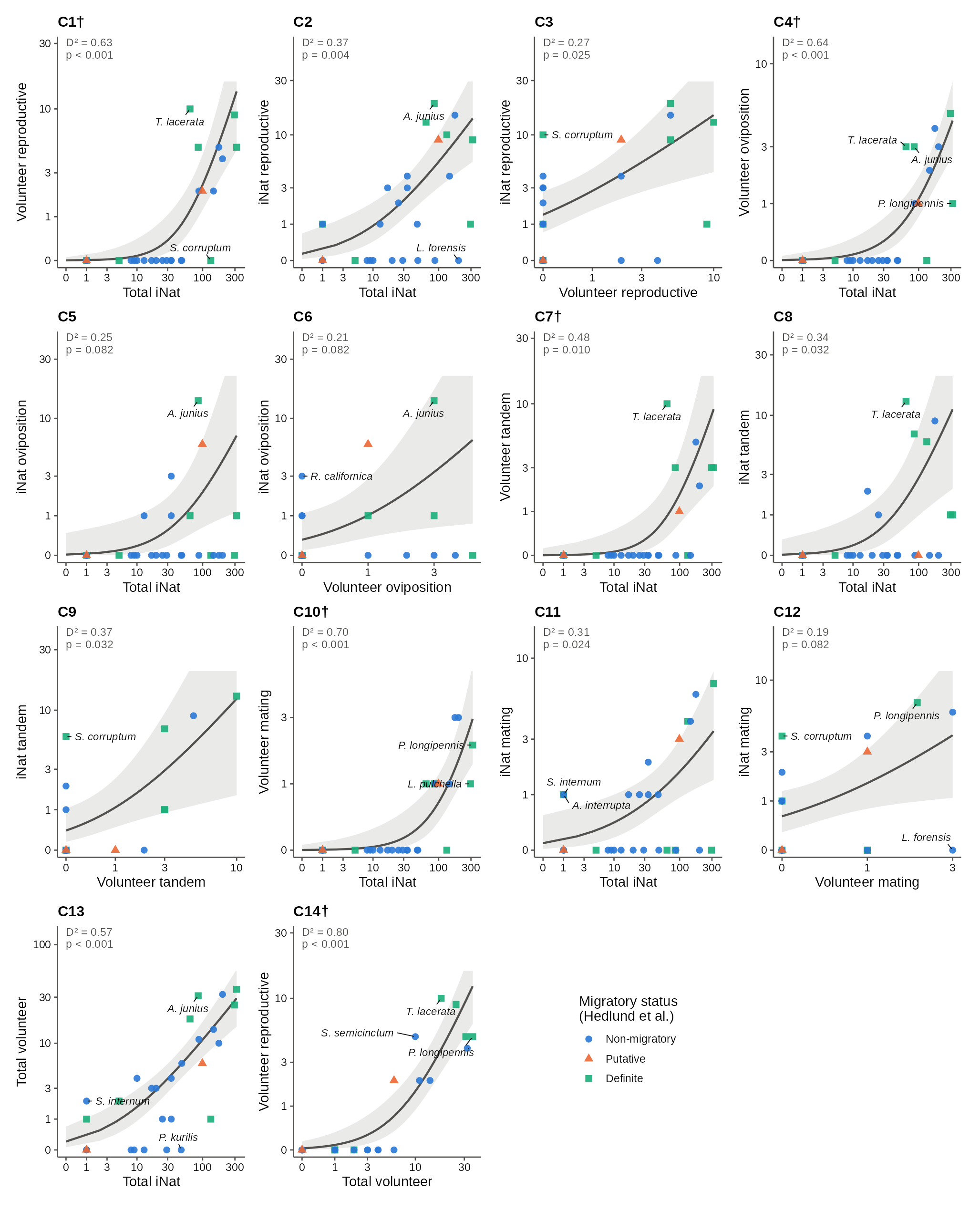

**Figure S2.** Associations between species-level observation counts (n = 28 species) for the 14 comparisons. Each panel shows a count GLM (log link; negative binomial or Poisson) of the y-axis count on log(x + 1) of the x-axis count. The fitted line is shown with its 95% confidence band, and both axes are on a log(x + 1) scale. Points are species, colored and shaped by migratory status (Hedlund et al., 2026). Labelled species are outliers (|studentized residual| > 2) or influential (Cook’s distance > 4/n). Each panel gives the deviance explained (D²) and the Holm-adjusted likelihood-ratio p-value. Comparisons: C1, volunteer reproductive vs. total iNaturalist observations; C2, iNaturalist reproductive vs. total iNaturalist observations; C3, iNaturalist reproductive vs. volunteer reproductive observations; C4–C6, C7–C9 and C10–C12, the same three comparisons for oviposition, tandem and mating, respectively; C13, total volunteer vs. total iNaturalist observations; C14, volunteer reproductive vs. total volunteer observations. Reproductive totals count each observation once per species. † Curvature test significant: volunteers rarely recorded behavior in less-observed species, which gives a threshold pattern; the linear model was retained.

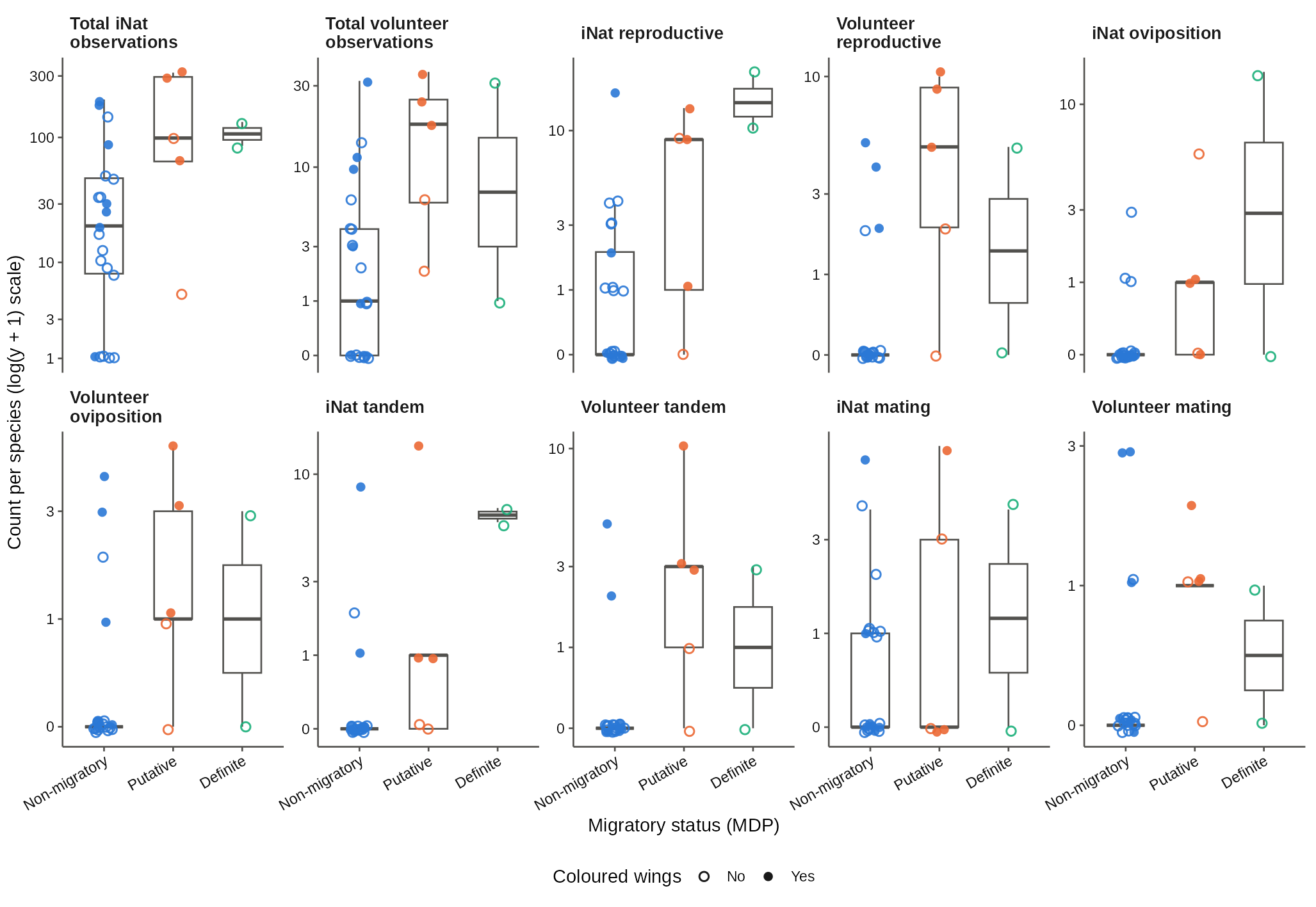

**Figure S3.** Observation counts per species (n = 28) by migratory status (Migratory Dragonfly Partnership) for each of the ten counts analyzed. Points are species, jittered horizontally; filled points have colored wings and open points do not. Boxes show the median and interquartile range, with whiskers to 1.5× the interquartile range. Counts are on a log(y + 1) scale, and y-axes differ among panels. Reproductive totals count each observation once per species.

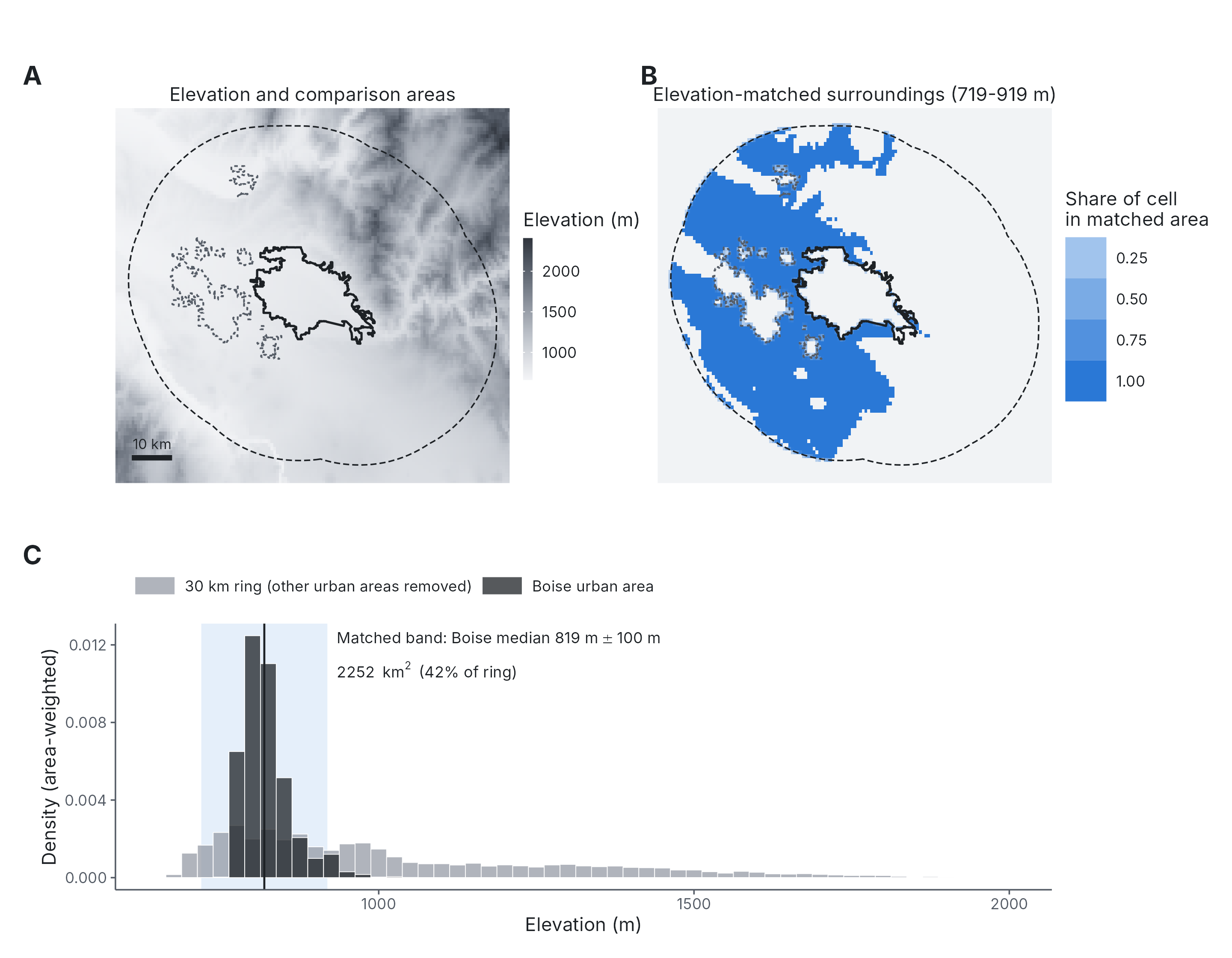

**Figure S4.** Comparison areas and elevation matching. (A) Elevation around Boise, with the Boise urban area (solid outline), the outer edge of the 30 km ring (dashed) and the excluded urban areas (dotted). Elevation from AWS Terrain Tiles (R package elevatr; c. 100 m resolution), averaged onto the 800 m PRISM grid. (B) Elevation-matched surroundings: ring cells between 719 and 919 m, within 100 m of Boise’s median elevation; shading shows the share of each 800 m cell inside the matched area. (C) Area-weighted elevation distributions of the Boise urban area (dark) and of the 30 km ring with other urban areas removed (grey). The shaded band is the matching band and the vertical line is Boise’s median elevation (819 m). The matched area covers 2,252 km² (42% of the ring). Scale bar: 10 km.

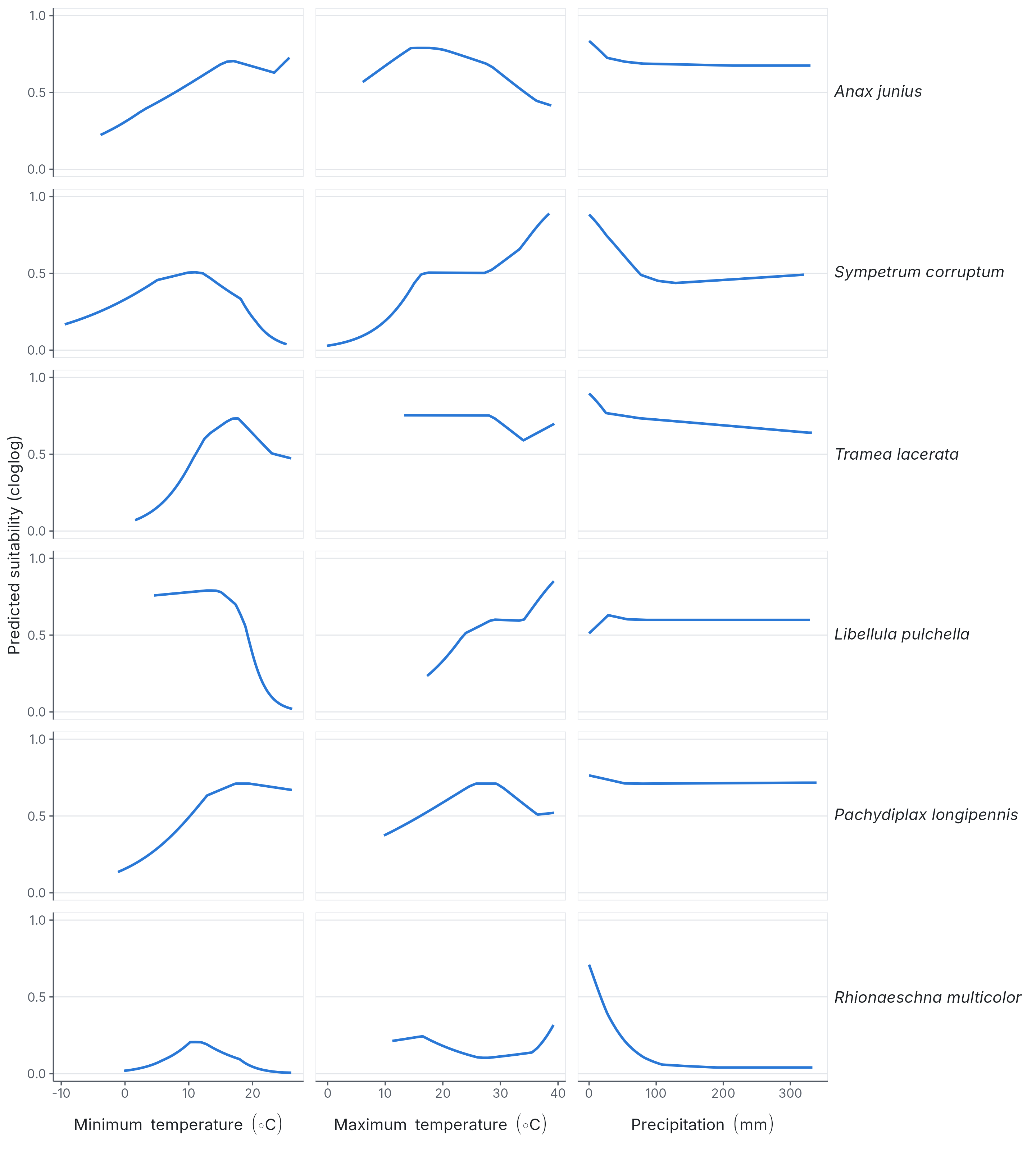

**Figure S5.** Response curves of the selected maxnet model for each species (LQH features, regularization multiplier 1). Each curve shows predicted suitability (cloglog) as one predictor varies across its 1st–99th percentile, with the other two held at their background medians. The percentiles are taken from thinned occurrences and background points together. Predictors are monthly PRISM minimum temperature, maximum temperature and total precipitation for the year-month of each record.

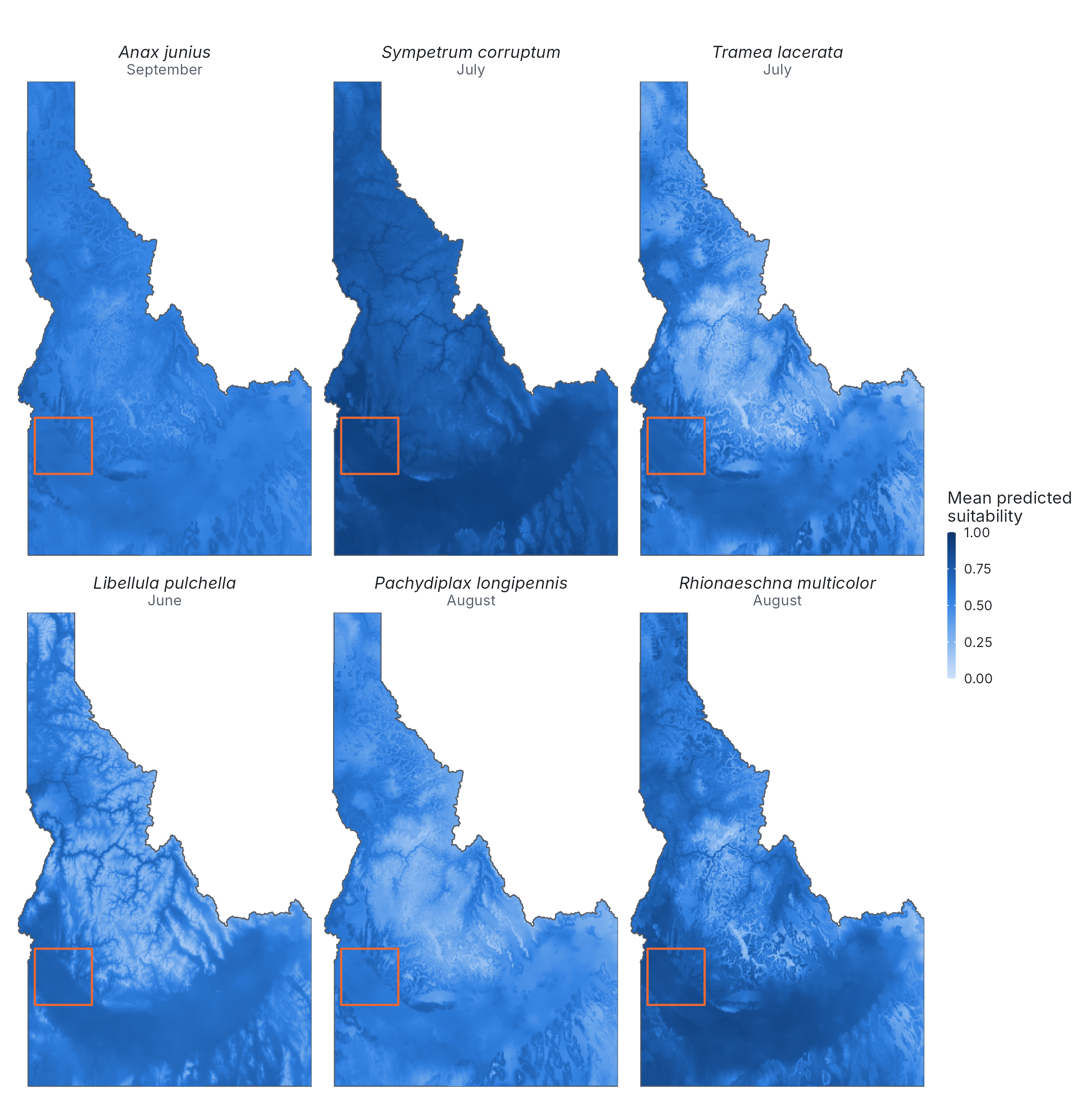

**Figure S6.** Long-term mean (1990–2025) predicted suitability across Idaho in each species’ peak month in Boise. The orange box marks the area shown in Figure 3B and Figure S7. For display, maps were aggregated to c. 1.6 km cells (2 × 2 cells of the 800 m grid).

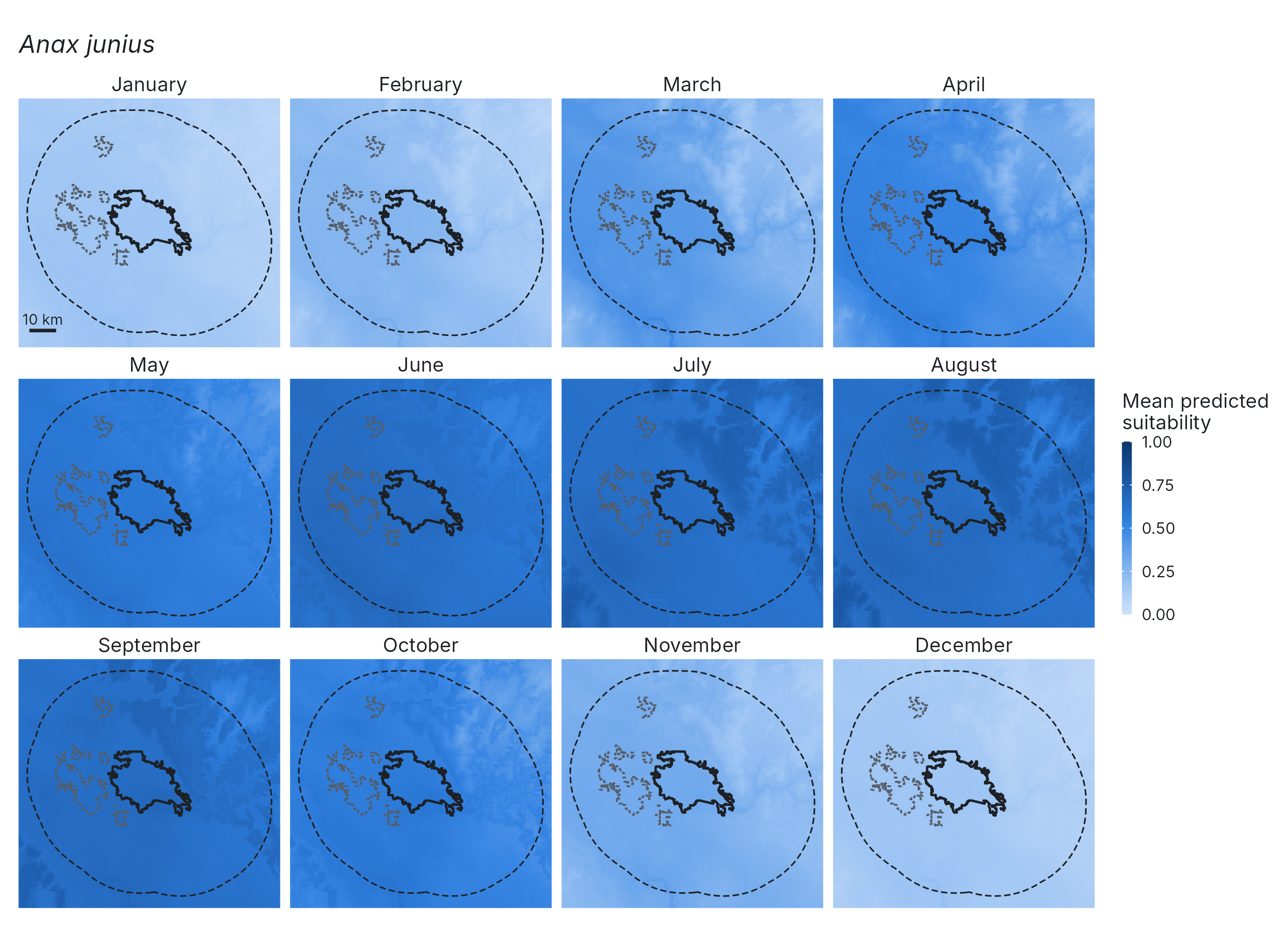

**Figure S7a.** *Anax junius*.

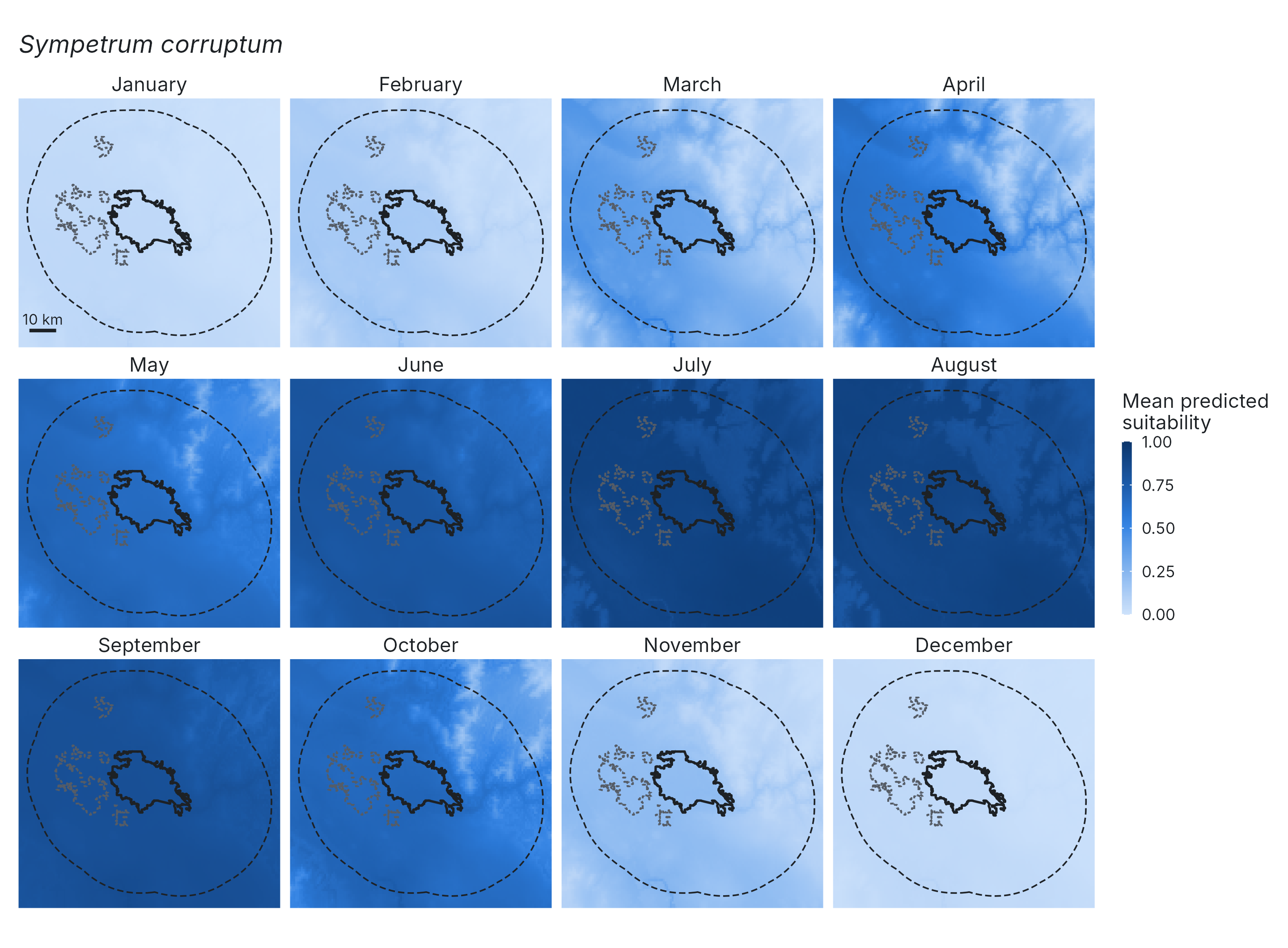

**Figure S7b.** *Sympetrum corruptum*.

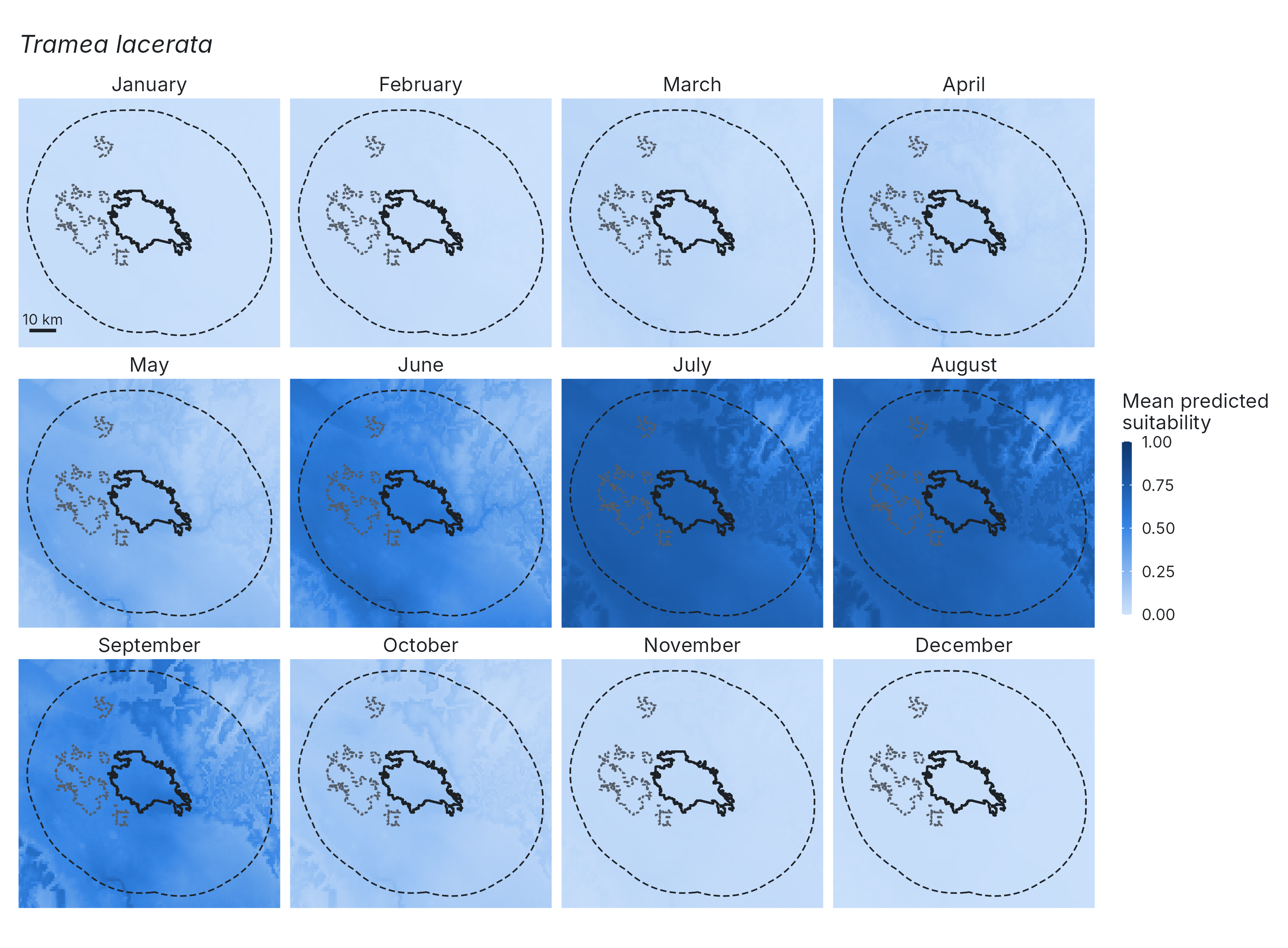

**Figure S7c.** *Tramea lacerata*.

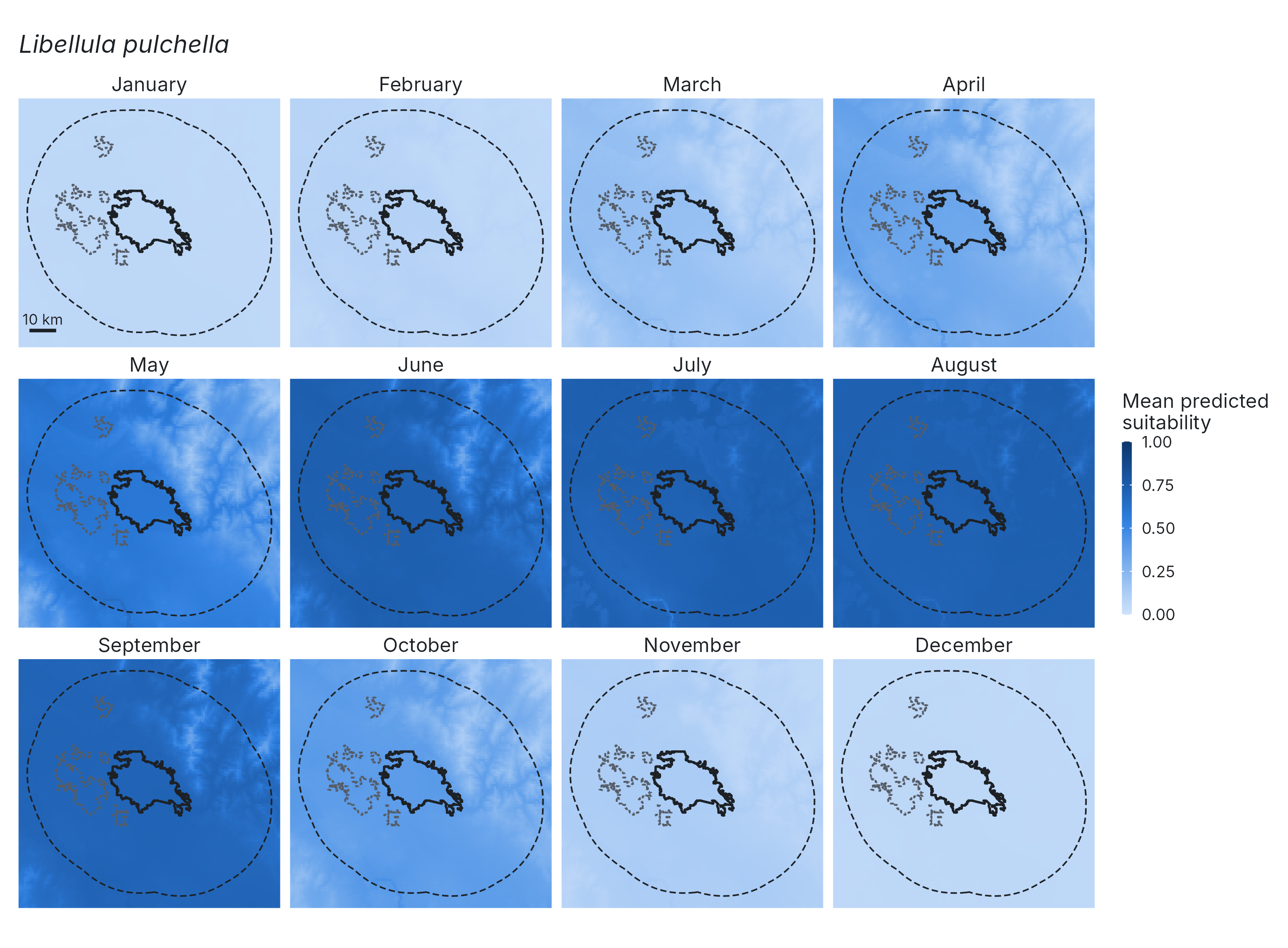

**Figure S7d.** *Libellula pulchella*.

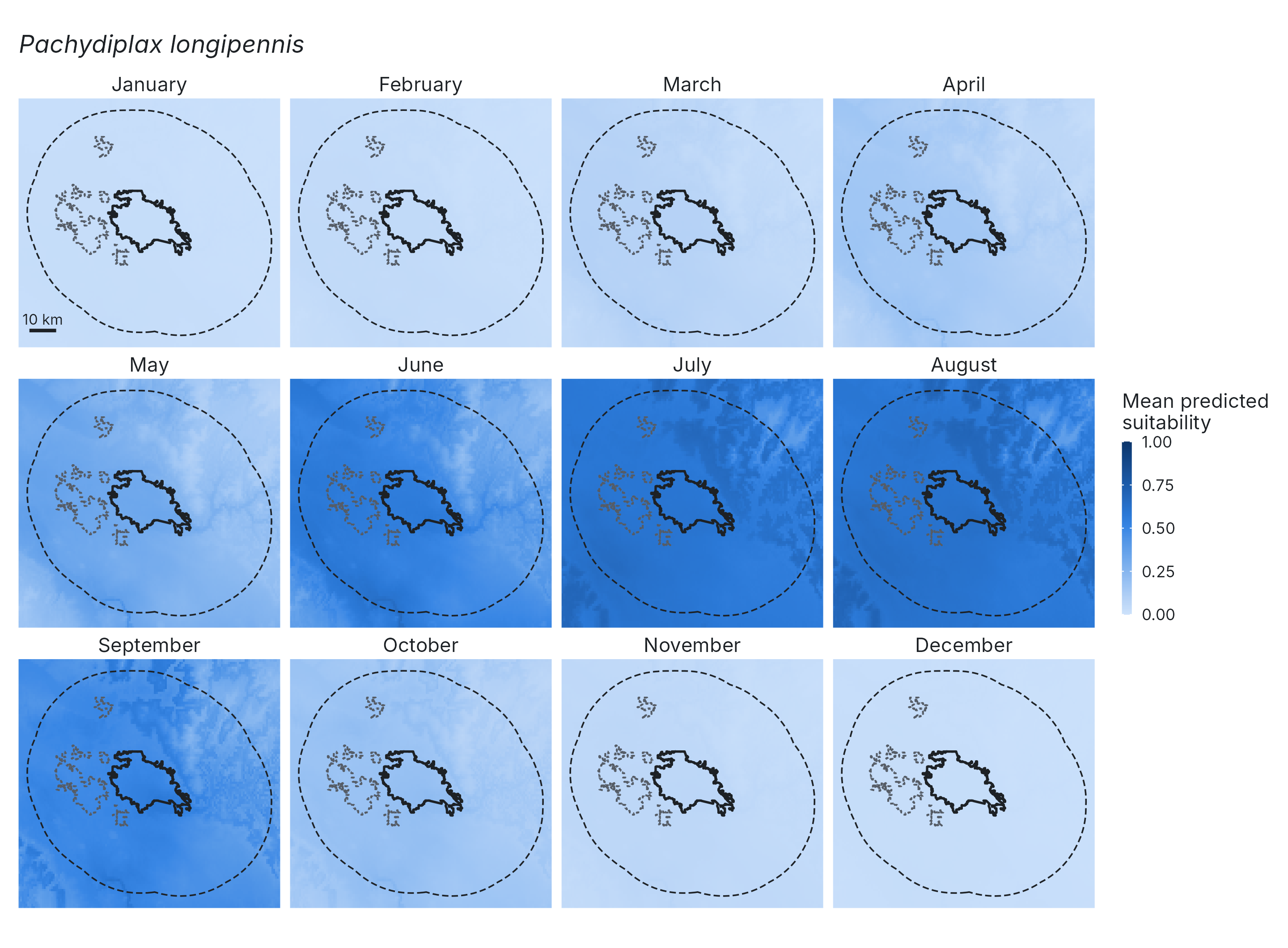

**Figure S7e.** *Pachydiplax longipennis*.

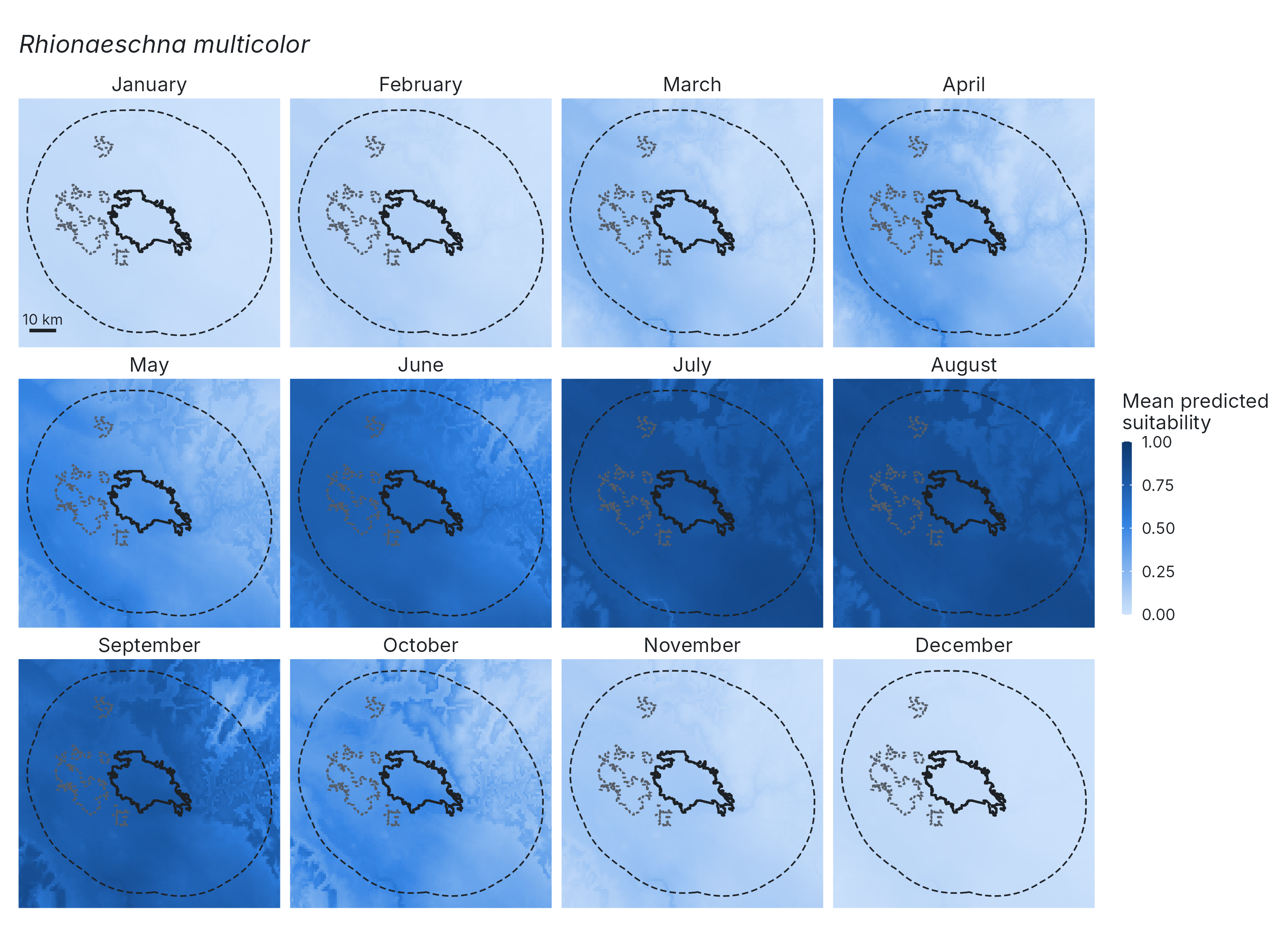

**Figure S7f.** *Rhionaeschna multicolor*.

**Figure S7.** Long-term mean (1990–2025) predicted suitability around Boise in each calendar month for (a) *Anax junius*, (b) *Sympetrum corruptum*, (c) *Tramea lacerata*, (d) *Libellula pulchella*, (e) *Pachydiplax longipennis* and (f) *Rhionaeschna multicolor*. Outlines are as in Figure 3B: Boise urban area solid, 30 km ring edge dashed, excluded urban areas dotted. All panels share a 0–1 color scale. Scale bar: 10 km.

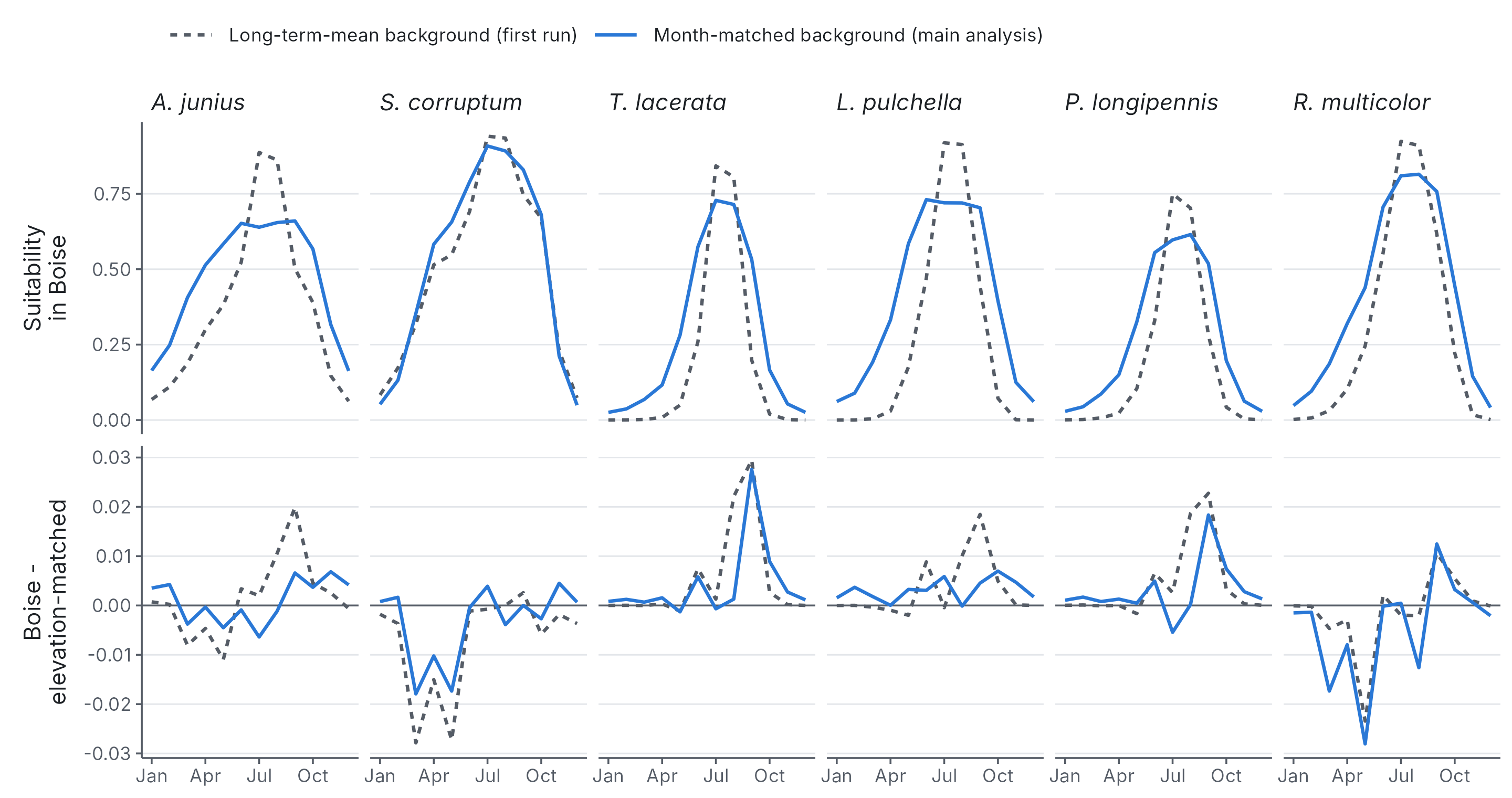

**Figure S8.** Sensitivity of the Boise comparison to the background design. Top row: mean predicted suitability in Boise by calendar month. Bottom row: Boise minus elevation-matched surroundings. Blue solid lines are the main analysis, in which each background point was given the climate of a year-month drawn in proportion to the species’ thinned records. Grey dashed lines are a run in which each background point was given its climate averaged over all 432 months of 1990–2025. Values are means across 36 years.
