## Supplemental Tables for "Sex in the City: Reproductive Behaviors of Dragonflies (Odonata: Anisoptera) across an Urban Landscape"

**Table S1.** Species included in the analyses, with authorities and common names (World Odonata List; Paulson et al., 2026), trait classifications and observation counts.

| **Family** | **Species (common name)** | **Hedlund** | **MDP** | **Flight** | **Color** | **Spots** | **iNat total** | **Vol. total** | **iNat repro.** | **Vol. repro.** | **iNat ovip.** | **Vol. ovip.** | **iNat tand.** | **Vol. tand.** | **iNat mat.** | **Vol. mat.** |
| --- | --- | --- | --- | --- | --- | --- | --- | --- | --- | --- | --- | --- | --- | --- | --- | --- |
| Aeshnidae | *Aeshna interrupta* Walker variable darner | Definite | Non-migratory | Flyer | No | No | 1 | 1 | 1 | 0 | 0 | 0 | 0 | 0 | 1 | 0 |
|  | *Aeshna palmata* Hagen paddle-tailed darner | Non-migratory | Non-migratory | Flyer | No | No | 34 | 1 | 3 | 0 | 1 | 0 | 0 | 0 | 2 | 0 |
|  | *Aeshna umbrosa* Walker shadow darner | Non-migratory | Non-migratory | Flyer | No | No | 13 | 0 | 1 | 0 | 1 | 0 | 0 | 0 | 0 | 0 |
|  | *Anax junius* (Drury) common green darner | Definite | Definite | Flyer | No | No | 86 | 31 | 19 | 5 | 14 | 3 | 7 | 3 | 0 | 1 |
|  | *Rhionaeschna californica* (Calvert) California darner | Non-migratory | Non-migratory | Flyer | No | No | 34 | 4 | 4 | 0 | 3 | 0 | 0 | 0 | 1 | 0 |
|  | *Rhionaeschna multicolor* (Hagen) blue-eyed darner | Putative | Possible migrant | Flyer | No | No | 99 | 6 | 9 | 2 | 6 | 1 | 0 | 1 | 3 | 1 |
| Cordulegastridae | *Zoraena dorsalis* (Hagen in Selys) Pacific spiketail | Non-migratory | Non-migratory | Flyer | No | No | 9 | 0 | 0 | 0 | 0 | 0 | 0 | 0 | 0 | 0 |
| Corduliidae | *Epitheca sp.* a baskettail | Putative | - | Flyer | Yes | Yes | 1 | 0 | 0 | 0 | 0 | 0 | 0 | 0 | 0 | 0 |
| Gomphidae | *Ophiogomphus occidentis* (Hagen) sinuous snaketail | Non-migratory | Non-migratory | Percher | No | No | 8 | 0 | 0 | 0 | 0 | 0 | 0 | 0 | 0 | 0 |
|  | *Ophiogomphus severus* Hagen in Hayden pale snaketail | Non-migratory | Non-migratory | Percher | No | No | 1 | 0 | 0 | 0 | 0 | 0 | 0 | 0 | 0 | 0 |
|  | *Phanogomphus kurilis* (Hagen in Selys) Pacific clubtail | Non-migratory | Non-migratory | Percher | No | No | 48 | 0 | 1 | 0 | 0 | 0 | 0 | 0 | 1 | 0 |
| Libellulidae | *Erythemis collocata* (Hagen) western pondhawk | Non-migratory | Non-migratory | Percher | No | No | 145 | 14 | 4 | 2 | 0 | 2 | 0 | 0 | 4 | 1 |
|  | *Leucorrhinia intacta* (Hagen) dot-tailed whiteface | Non-migratory | Non-migratory | Percher | Yes | No | 25 | 1 | 2 | 0 | 0 | 0 | 1 | 0 | 1 | 0 |
|  | *Libellula forensis* Hagen eight-spotted skimmer | Non-migratory | Non-migratory | Percher | Yes | Yes | 197 | 32 | 0 | 4 | 0 | 3 | 0 | 2 | 0 | 3 |
|  | *Libellula pulchella* Drury twelve-spotted skimmer | Definite | Putative | Intermediate | Yes | Yes | 295 | 25 | 1 | 9 | 0 | 5 | 1 | 3 | 0 | 1 |
|  | *Libellula quadrimaculata* Linnaeus four-spotted skimmer | Non-migratory | Non-migratory | Percher | Yes | Yes | 20 | 3 | 0 | 0 | 0 | 0 | 0 | 0 | 0 | 0 |
|  | *Libellula saturata* Uhler flame skimmer | Non-migratory | Non-migratory | Percher | Yes | Yes | 29 | 0 | 0 | 0 | 0 | 0 | 0 | 0 | 0 | 0 |
|  | *Pachydiplax longipennis* (Burmeister) blue dasher | Definite | Putative | Percher | Yes | No | 318 | 36 | 9 | 5 | 1 | 1 | 1 | 3 | 7 | 2 |
|  | *Plathemis lydia* (Drury) common whitetail | Non-migratory | Non-migratory | Percher | Yes | Yes | 88 | 11 | 0 | 2 | 0 | 1 | 0 | 0 | 0 | 1 |
|  | *Sympetrum corruptum* (Hagen) variegated meadowhawk | Definite | Definite | Percher | No | No | 132 | 1 | 10 | 0 | 0 | 0 | 6 | 0 | 4 | 0 |
|  | *Sympetrum costiferum* (Hagen) saffron-winged meadowhawk | Non-migratory | Non-migratory | Percher | No | No | 17 | 3 | 3 | 0 | 0 | 0 | 2 | 0 | 1 | 0 |
|  | *Sympetrum internum* Montgomery cherry-faced meadowhawk | Non-migratory | Non-migratory | Percher | No | No | 1 | 2 | 1 | 0 | 0 | 0 | 0 | 0 | 1 | 0 |
|  | *Sympetrum obtrusum* (Hagen) white-faced meadowhawk | Non-migratory | Non-migratory | Percher | No | No | 10 | 4 | 0 | 0 | 0 | 0 | 0 | 0 | 0 | 0 |
|  | *Sympetrum pallipes* (Hagen in Hayden) striped meadowhawk | Non-migratory | Non-migratory | Percher | No | No | 49 | 6 | 0 | 0 | 0 | 0 | 0 | 0 | 0 | 0 |
|  | *Sympetrum semicinctum* (Say) band-winged meadowhawk | Non-migratory | Non-migratory | Percher | Yes | Yes | 174 | 10 | 15 | 5 | 0 | 4 | 9 | 5 | 6 | 3 |
|  | *Sympetrum vicinum* (Hagen) autumn meadowhawk | Definite | Putative | Percher | No | No | 5 | 2 | 0 | 0 | 0 | 0 | 0 | 0 | 0 | 0 |
|  | *Tramea lacerata* Hagen black saddlebags | Definite | Putative | Flyer | Yes | Yes | 65 | 18 | 13 | 10 | 1 | 3 | 13 | 10 | 0 | 1 |
| Macromiidae | *Macromia magnifica* Selys western river cruiser | Non-migratory | Non-migratory | Flyer | No | No | 1 | 0 | 0 | 0 | 0 | 0 | 0 | 0 | 0 | 0 |

MDP ‘–’ = not listed (coded Non-migratory in the analyses). Reproductive totals count each iNat record or volunteer survey response once per species, whatever behaviors it shows.

**Table S2.** Assumption checks for the best trait model of each observation count.

| **Response** | **Family** | **SW p** | **Dispersion** | **Zeros obs/exp** | **Zero p** | **Fligner p** | **GVIF** | **Family RE var.** | **Signs kept** | **Outliers / influential** | **Single-species class** |
| --- | --- | --- | --- | --- | --- | --- | --- | --- | --- | --- | --- |
| Total iNat observations | NB (θ = 0.80) | 0.419 | 0.88 | 0 / 1.6 | 1.000 | 0.971 | – | family fixed | – | *S. internum, Z. dorsalis, P. kurilis* | *–* |
| Total volunteer observations | NB (θ = 0.89) | 0.892 | 0.89 | 8 / 8.9 | 0.800 | 0.693 | 1.18 | family fixed | – | *A. junius, A. umbrosa, L. saturata* | *–* |
| iNat reproductive | NB (θ = 0.57) | 0.440 | 1.44 | 12 / 11.2 | 0.459 | 0.557 | – | 0.00 | yes | *S. semicinctum* | *–* |
| Volunteer reproductive | NB (θ = 1.20) | 0.894 | 1.35 | 19 / 16.5 | 0.173 | 0.186 | 1.16 | 0.00 | yes | *S. corruptum, E. collocata, A. junius, S. semicinctum* | *–* |
| iNat oviposition | Poisson | 0.642 | 0.58 | 21 / 21.1 | 0.666 | 0.166 | 1.03 | 0.00 | yes | *R. californica, A. junius, P. longipennis, T. lacerata, R. multicolor* | *–* |
| Volunteer oviposition | NB (θ = 3.13) | 0.706 | 1.46 | 19 / 17.5 | 0.316 | 0.511 | 1.16 | 0.00 | yes | *S. corruptum, E. collocata, A. junius, S. semicinctum* | *–* |
| iNat tandem | NB (θ = 0.44) | 0.600 | 1.57 | 20 / 19.5 | 0.504 | 0.359 | 1.15 | 0.00 | yes | *S. semicinctum, S. costiferum, T. lacerata* | *–* |
| Volunteer tandem | NB (θ = 1.07) | 0.634 | 0.77 | 21 / 20.0 | 0.385 | 0.581 | 1.46 | 0.00 | yes | *S. corruptum, S. semicinctum, A. junius, R. multicolor* | *–* |
| iNat mating | NB (θ = 0.43) | 0.699 | 0.88 | 16 / 16.0 | 0.595 | – | – | – | – | *P. longipennis, S. semicinctum* | *–* |
| Volunteer mating | Poisson | 0.474 | 1.03 | 19 / 18.6 | 0.530 | 0.211 | – | 0.00 | yes | *L. forensis, S. semicinctum* | *–* |

SW p, Shapiro–Wilk test of randomized quantile residuals; Dispersion, Pearson χ²/df; Zero p, proportion of 2,000 simulated datasets with at least the observed number of zeros; Fligner p, smallest Fligner–Killeen p across the model's predictors; GVIF, max GVIF^(1/(2df)); Family RE var., variance of a family-level random intercept when the best model is refitted as a GLMM (very large values arise where whole families have zero counts, e.g. no volunteer records of Gomphidae, Cordulegastridae, Corduliidae or Macromiidae); Signs kept, whether all estimable trait effects keep their sign.

**Table S3.** AICc model selection for behavior counts adjusted for observation frequency: log(total observations + 1) from the same source is included in every candidate model.

| **Response** |  | **Model** | **Family** | **k** | **logLik** | **AICc** | **ΔAICc** | **w** | **D²** |
| --- | --- | --- | --- | --- | --- | --- | --- | --- | --- |
| iNat reproductive | • | Obs. frequency + Flight style (2 classes) | NB | 4 | -52.8 | 115.4 | 0.00 | 0.21 | 0.51 |
|  |  | Obs. frequency + Flight style (3 classes) | NB | 5 | -51.4 | 115.6 | 0.15 | 0.19 | 0.58 |
|  |  | Observation frequency only | NB | 3 | -55.7 | 118.3 | 2.91 | 0.05 | 0.37 |
| Volunteer reproductive | • | Obs. frequency + Flight style (2 classes) + Wing spots/bands | Poisson | 4 | -23.0 | 55.7 | 0.00 | 0.15 | 0.86 |
|  |  | Obs. frequency + Migratory (Hedlund et al.) + Wing spots/bands | Poisson | 5 | -21.7 | 56.1 | 0.46 | 0.12 | 0.88 |
|  |  | Obs. frequency + Migratory (MDP) + Wing spots/bands | Poisson | 5 | -21.9 | 56.6 | 0.89 | 0.09 | 0.88 |
|  |  | Obs. frequency + Flight style (2 classes) + Wing color | Poisson | 4 | -23.8 | 57.4 | 1.75 | 0.06 | 0.84 |
|  |  | Obs. frequency + Flight style (3 classes) + Wing color | Poisson | 5 | -22.4 | 57.5 | 1.87 | 0.06 | 0.87 |
|  |  | Obs. frequency + Wing spots/bands | Poisson | 3 | -25.3 | 57.6 | 1.93 | 0.06 | 0.82 |
|  |  | Obs. frequency + Flight style (3 classes) + Wing spots/bands | Poisson | 5 | -22.4 | 57.6 | 1.94 | 0.06 | 0.87 |
|  |  | Observation frequency only | NB | 3 | -26.9 | 60.7 | 5.02 | 0.01 | 0.80 |
| iNat oviposition | • | Obs. frequency + Flight style (2 classes) + Wing spots/bands | Poisson | 4 | -14.0 | 37.7 | 0.00 | 0.15 | 0.92 |
|  |  | Obs. frequency + Migratory (Hedlund et al.) + Flight style (2 classes) + Wing spots/bands | Poisson | 6 | -11.1 | 38.3 | 0.61 | 0.11 | 0.97 |
|  |  | Obs. frequency + Migratory (MDP) + Flight style (2 classes) | Poisson | 5 | -13.0 | 38.7 | 1.01 | 0.09 | 0.93 |
|  |  | Obs. frequency + Flight style (2 classes) + Wing color + Wing spots/bands | Poisson | 5 | -13.3 | 39.3 | 1.60 | 0.07 | 0.93 |
|  |  | Obs. frequency + Migratory (MDP) + Flight style (2 classes) + Wing spots/bands | Poisson | 6 | -11.8 | 39.5 | 1.85 | 0.06 | 0.96 |
|  |  | Observation frequency only | NB | 3 | -27.9 | 62.7 | 25.05 | 0.00 | 0.25 |
| Volunteer oviposition | • | Obs. frequency + Wing spots/bands | Poisson | 3 | -18.6 | 44.2 | 0.00 | 0.29 | 0.78 |
|  |  | Obs. frequency + Wing color + Wing spots/bands | Poisson | 4 | -17.7 | 45.2 | 1.00 | 0.18 | 0.81 |
|  |  | Observation frequency only | Poisson | 2 | -21.1 | 46.7 | 2.53 | 0.08 | 0.70 |
| iNat tandem | • | Observation frequency only | NB | 3 | -33.0 | 73.1 | 0.00 | 0.25 | 0.34 |
|  |  | Obs. frequency + Flight style (2 classes) | NB | 4 | -32.5 | 74.7 | 1.63 | 0.11 | 0.39 |
|  |  | Obs. frequency + Migratory (Hedlund et al.) | NB | 5 | -31.0 | 74.7 | 1.67 | 0.11 | 0.49 |
| Volunteer tandem | • | Obs. frequency + Flight style (2 classes) + Wing color | Poisson | 4 | -18.6 | 47.0 | 0.00 | 0.38 | 0.82 |
|  |  | Observation frequency only | NB | 3 | -23.2 | 53.5 | 6.55 | 0.01 | 0.68 |
| iNat mating |  | Obs. frequency + Wing spots/bands | Poisson | 3 | -34.8 | 76.7 | 0.00 | 0.18 | 0.47 |
|  |  | Obs. frequency + Flight style (3 classes) + Wing spots/bands | Poisson | 5 | -32.5 | 77.7 | 0.99 | 0.11 | 0.54 |
|  | •‡ | Obs. frequency + Wing spots/bands | NB | 4 | -34.1 | 77.9 | 1.28 | 0.10 | 0.42 |
|  |  | Observation frequency only | NB | 3 | -35.9 | 78.8 | 2.10 | 0.06 | 0.31 |
| Volunteer mating | • | Observation frequency only | Poisson | 2 | -15.9 | 36.4 | 0.00 | 0.20 | 0.68 |
|  |  | Obs. frequency + Wing spots/bands | Poisson | 3 | -15.0 | 37.1 | 0.68 | 0.14 | 0.73 |
|  |  | Obs. frequency + Wing color | Poisson | 3 | -15.3 | 37.5 | 1.18 | 0.11 | 0.72 |
|  |  | Obs. frequency + Migratory (Hedlund et al.) | Poisson | 4 | -14.1 | 38.0 | 1.64 | 0.09 | 0.78 |

• Retained (best) model. ‡ The AICc-best Poisson model failed its distributional checks, so the lowest-AICc negative binomial model was retained. NB, negative binomial; k, number of estimated parameters (including θ for NB); w, Akaike weight; D², deviance explained.

**Table S4.** Trait effects in the best observation-frequency-adjusted models (IRR, 95% profile CI).

| **Response** | **Predictor** | **Level** | **n** | **IRR (95% CI)** | **LRT χ² (df)** | **p** |
| --- | --- | --- | --- | --- | --- | --- |
| iNat reproductive | Flight style (2 classes) | Flyer vs Percher | 10 | 3.50 (1.40, 9.51) | 5.65 (1) | 0.017 |
| Volunteer reproductive | Flight style (2 classes) | Flyer vs Percher | 10 | 2.05 (1.07, 3.82) | 4.67 (1) | 0.031 |
|  | Wing spots/bands | Yes vs No | 8 | 2.91 (1.54, 5.78) | 11.06 (1) | <0.001 |
| iNat oviposition | Flight style (2 classes) | Flyer vs Percher | 10 | 133.60 (23.74, 2579.29) | 61.91 (1) | <0.001 |
|  | Wing spots/bands | Yes vs No | 8 | 0.15 (0.01, 0.71) | 6.42 (1) | 0.011 |
| Volunteer oviposition | Wing spots/bands | Yes vs No | 8 | 2.66 (1.13, 6.95) | 5.05 (1) | 0.025 |
| iNat tandem | Observation frequency only | – |  | – | – | – |
| Volunteer tandem | Flight style (2 classes) | Flyer vs Percher | 10 | 5.60 (2.47, 12.72) | 16.08 (1) | <0.001 |
|  | Wing color | Yes vs No | 10 | 7.40 (2.63, 26.31) | 15.72 (1) | <0.001 |
| iNat mating | Wing spots/bands | Yes vs No | 8 | 0.31 (0.09, 0.95) | 3.55 (1) | 0.060 |
| Volunteer mating | Observation frequency only | – |  | – | – | – |

IRRs are relative to the reference class at a given observation frequency.

**Table S5.** Sensitivity of the best raw-count model to the coding of species not listed in MDP (Epitheca sp.): dropped, or given their own class.

| **Response** | **Primary (– = Non-migratory)** | **– species dropped** | **– as own class** | **Agreement** |
| --- | --- | --- | --- | --- |
| Total iNat observations | Family | Wing color | Migratory (MDP) + Wing color | differs |
| Total volunteer observations | Wing color + Family | Wing color + Family | Wing color + Family | same |
| iNat reproductive | Migratory (MDP) | Migratory (MDP) | Family | differs |
| Volunteer reproductive | Migratory (MDP) + Wing color | Migratory (MDP) + Wing color + Wing spots/bands | Migratory (MDP) + Wing color + Wing spots/bands | differs |
| iNat oviposition | Migratory (MDP) + Flight style (2 classes) + Wing spots/bands | Migratory (MDP) + Flight style (2 classes) + Wing spots/bands | Migratory (MDP) + Flight style (2 classes) + Wing spots/bands | same |
| Volunteer oviposition | Migratory (MDP) + Wing spots/bands | Migratory (MDP) + Wing spots/bands | Migratory (MDP) + Wing spots/bands | same |
| iNat tandem | Migratory (MDP) + Wing color | Migratory (MDP) + Wing color | Migratory (MDP) + Wing color | same |
| Volunteer tandem | Migratory (MDP) + Wing color | Migratory (MDP) + Flight style (2 classes) + Wing color | Migratory (MDP) + Flight style (2 classes) + Wing color | differs |
| iNat mating | Null (intercept only) | Null (intercept only) | Null (intercept only) | same |
| Volunteer mating | Wing color | Wing color | Wing color | same |

**Table S6.** Fit and assumption checks for all 32 models: the 14 count comparisons (C1–C14), the best raw-count trait models (T1–T10) and the best observation-frequency-adjusted trait models (A1–A8). n = 28 species for every model.

| **Set** | **ID** | **Model** | **Family** | **D²** | **SW p** | **Dispersion** | **Zeros obs. (exp.; 95% sim.)** | **Zero p** | **Form / homog. p** | **Max Cook's D** | **Flagged** |
| --- | --- | --- | --- | --- | --- | --- | --- | --- | --- | --- | --- |
| Count comparisons | C1 | Volunteer reproductive obs. ~ Total iNat obs. | NB (θ = 0.89) | 0.63 | 0.112 | 1.28 | 19 (17.9; 14–22) | 0.360 | 0.024 | 1.00 | 2 |
|  | C2 | iNat reproductive obs. ~ Total iNat obs. | NB (θ = 0.73) | 0.37 | 0.508 | 0.76 | 12 (11.6; 7–16) | 0.513 | 0.884 | 0.16 | 2 |
|  | C3 | iNat reproductive obs. ~ Volunteer reproductive obs. | NB (θ = 0.60) | 0.27 | 0.609 | 1.03 | 12 (11.2; 6–16) | 0.456 | 0.658 | 0.45 | 1 |
|  | C4 | Volunteer oviposition obs. ~ Total iNat obs. | Poisson | 0.64 | 0.490 | 0.83 | 19 (17.9; 14–21) | 0.391 | 0.021 | 1.23 | 3 |
|  | C5 | iNat oviposition obs. ~ Total iNat obs. | NB (θ = 0.19) | 0.25 | 0.579 | 0.59 | 21 (21.4; 17–25) | 0.670 | 0.061 | 0.28 | 1 |
|  | C6 | iNat oviposition obs. ~ Volunteer oviposition obs. | NB (θ = 0.20) | 0.21 | 0.687 | 0.74 | 21 (21.1; 17–25) | 0.607 | 0.218 | 0.25 | 2 |
|  | C7 | Volunteer tandem obs. ~ Total iNat obs. | NB (θ = 0.38) | 0.48 | 0.330 | 1.28 | 21 (20.8; 17–25) | 0.566 | 0.046 | 0.99 | 1 |
|  | C8 | iNat tandem obs. ~ Total iNat obs. | NB (θ = 0.25) | 0.34 | 0.539 | 0.69 | 20 (20.0; 16–24) | 0.603 | 0.109 | 0.28 | 1 |
|  | C9 | iNat tandem obs. ~ Volunteer tandem obs. | NB (θ = 0.36) | 0.37 | 0.436 | 1.75 | 20 (19.2; 14–24) | 0.478 | 0.658 | 0.91 | 1 |
|  | C10 | Volunteer mating obs. ~ Total iNat obs. | Poisson | 0.70 | 0.493 | 0.35 | 19 (20.4; 17–24) | 0.868 | 0.045 | 0.35 | 2 |
|  | C11 | iNat mating obs. ~ Total iNat obs. | NB (θ = 1.15) | 0.31 | 0.181 | 0.84 | 16 (15.4; 11–20) | 0.487 | 0.194 | 0.20 | 2 |
|  | C12 | iNat mating obs. ~ Volunteer mating obs. | NB (θ = 0.72) | 0.19 | 0.486 | 0.84 | 16 (15.6; 11–20) | 0.516 | 0.875 | 0.27 | 3 |
|  | C13 | Total volunteer obs. ~ Total iNat obs. | NB (θ = 1.42) | 0.57 | 0.644 | 0.99 | 8 (7.1; 3–11) | 0.410 | 0.378 | 0.25 | 3 |
|  | C14 | Volunteer reproductive obs. ~ Total volunteer obs. | NB (θ = 2.69) | 0.80 | 0.628 | 0.60 | 19 (18.1; 15–21) | 0.392 | <0.001 | 0.23 | 3 |
| Trait models (raw) | T1 | Total iNat observations ~ Family | NB (θ = 0.80) | 0.31 | 0.424 | 0.88 | 0 (1.5; 0–4) | 1.000 | 0.971 | 0.34 | 3 |
|  | T2 | Total volunteer observations ~ Color + Family | NB (θ = 0.89) | 0.55 | 0.930 | 0.89 | 8 (9.0; 6–12) | 0.821 | 0.693 | 0.42 | 3 |
|  | T3 | iNat reproductive ~ GEODE | NB (θ = 0.57) | 0.24 | 0.411 | 1.44 | 12 (11.1; 6–16) | 0.439 | 0.557 | 0.47 | 1 |
|  | T4 | Volunteer reproductive ~ GEODE + Color | NB (θ = 1.20) | 0.55 | 0.632 | 1.35 | 19 (16.6; 12–21) | 0.176 | 0.186 | 0.40 | 4 |
|  | T5 | iNat oviposition ~ GEODE + Flight + Spots | Poisson | 0.88 | 0.642 | 0.58 | 21 (21.1; 18–24) | 0.653 | 0.166 | 7.58 | 5 |
|  | T6 | Volunteer oviposition ~ GEODE + Spots | NB (θ = 3.13) | 0.47 | 0.600 | 1.46 | 19 (17.5; 13–21) | 0.320 | 0.511 | 0.51 | 4 |
|  | T7 | iNat tandem ~ GEODE + Color | NB (θ = 0.44) | 0.48 | 0.706 | 1.57 | 20 (19.5; 15–24) | 0.513 | 0.359 | 0.43 | 3 |
|  | T8 | Volunteer tandem ~ GEODE + Color | NB (θ = 1.07) | 0.64 | 0.629 | 0.77 | 21 (20.0; 16–23) | 0.401 | 0.581 | 0.47 | 4 |
|  | T9 | iNat mating ~ null | NB (θ = 0.43) | 0.00 | 0.716 | 0.88 | 16 (16.0; 11–21) | 0.571 | – | 0.31 | 2 |
|  | T10 | Volunteer mating ~ Color | Poisson | 0.30 | 0.487 | 1.03 | 19 (18.6; 14–23) | 0.523 | 0.211 | 0.20 | 2 |
| Trait models (adjusted) | A1 | iNat reproductive ~ log obs. + Flight | NB (θ = 1.25) | 0.51 | 0.387 | 1.12 | 12 (11.0; 7–15) | 0.399 | 0.159 | 0.16 | 4 |
|  | A2 | Volunteer reproductive ~ log obs. + Flight + Spots | Poisson | 0.86 | 0.604 | 0.61 | 19 (17.6; 14–21) | 0.295 | 0.405 | 1.33 | 5 |
|  | A3 | iNat oviposition ~ log obs. + Flight + Spots | Poisson | 0.92 | 0.517 | 0.33 | 21 (22.1; 20–24) | 0.892 | 0.454 | 1.86 | 4 |
|  | A4 | Volunteer oviposition ~ log obs. + Spots | Poisson | 0.78 | 0.498 | 0.52 | 19 (19.0; 16–22) | 0.639 | 0.829 | 0.53 | 3 |
|  | A5 | iNat tandem ~ log obs. | NB (θ = 0.25) | 0.34 | 0.640 | 0.69 | 20 (20.0; 16–24) | 0.600 | – | 0.28 | 1 |
|  | A6 | Volunteer tandem ~ log obs. + Flight + Color | Poisson | 0.82 | 0.155 | 0.92 | 21 (19.9; 17–23) | 0.370 | 0.656 | 0.70 | 2 |
|  | A7 | iNat mating ~ log obs. + Spots | NB (θ = 2.26) | 0.42 | 0.396 | 1.37 | 16 (15.1; 11–19) | 0.426 | 0.863 | 1.37 | 4 |
|  | A8 | Volunteer mating ~ log obs. | Poisson | 0.68 | 0.594 | 0.58 | 19 (20.4; 17–24) | 0.887 | – | 0.36 | 1 |

NB, negative binomial. D², deviance explained. SW p, Shapiro–Wilk test of randomized quantile residuals (median of 25 randomizations). Dispersion, Pearson χ²/df. Zeros: observed (expected; 95% range of 2,000 simulations); Zero p, proportion of simulations with at least the observed zeros. Form / homog. p: quadratic-term LRT (comparisons) or smallest Fligner–Killeen p across trait classes (trait models). Flagged: |studentized residual| > 2 or Cook's D > 4/n. log obs., log(total observations + 1) from the same source.

**Table S7.** Occurrence data, selected model settings and evaluation statistics for the six maxnet models. Evaluation statistics are means ± SD across five random cross-validation folds.

| Species | Records | Records after thinning | Year-months | Features, RM | Coef. | AUC (validation) | AUC diff | OR10 | CBI (validation) | Boyce (all data) | Threshold | Null AUC (mean) | z | p |
| --- | --- | --- | --- | --- | --- | --- | --- | --- | --- | --- | --- | --- | --- | --- |
| *Anax junius* | 23,885 | 14,074 | 187 | LQH, 1 | 23 | 0.608 ± 0.008 | 0.008 | 0.100 ± 0.008 | 0.986 ± 0.006 | 0.997 | 0.504 | 0.599 | 7.6 | < 0.001 |
| *Sympetrum corruptum* | 18,806 | 9,722 | 189 | LQH, 1 | 26 | 0.773 ± 0.005 | 0.005 | 0.100 ± 0.007 | 1.000 ± 0.000 | 1.000 | 0.359 | 0.755 | 14.2 | < 0.001 |
| *Tramea lacerata* | 9,187 | 5,923 | 123 | LQH, 1 | 16 | 0.644 ± 0.009 | 0.009 | 0.100 ± 0.006 | 0.993 ± 0.004 | 1.000 | 0.441 | 0.605 | 5.1 | < 0.001 |
| *Libellula pulchella* | 11,406 | 7,049 | 114 | LQH, 1 | 19 | 0.720 ± 0.004 | 0.004 | 0.100 ± 0.008 | 0.963 ± 0.035 | 0.995 | 0.425 | 0.705 | 5.0 | < 0.001 |
| *Pachydiplax longipennis* | 61,075 | 24,804 | 193 | LQH, 1 | 13 | 0.629 ± 0.004 | 0.003 | 0.101 ± 0.002 | 0.994 ± 0.003 | 0.998 | 0.532 | 0.625 | 6.2 | < 0.001 |
| *Rhionaeschna multicolor* | 4,714 | 2,886 | 114 | LQH, 1 | 25 | 0.875 ± 0.009 | 0.010 | 0.101 ± 0.016 | 0.994 ± 0.004 | 1.000 | 0.184 | 0.829 | 4.4 | < 0.001 |

Records: GBIF records (≤ 1 km coordinate uncertainty, 1990–2025) after removing exact duplicates. Records after thinning: records used to fit the model, after thinning to 20 km within each year-month, dropping year-months with < 5 records before or after thinning, and dropping records without PRISM values. Background: 50,000 points per species, each assigned a year-month in proportion to the species' thinned records. Features: L = linear, Q = quadratic, H = hinge; RM = regularization multiplier. The selected setting had the highest mean validation AUC (ties broken by lowest OR10) and also the lowest AICc (Akaike weight = 1) for every species.

AUC diff: training minus validation AUC. OR10: omission rate of held-out records at the 10th-percentile training-presence threshold (expected 0.10). CBI: continuous Boyce index on held-out folds; Boyce (all data): Boyce index of the final model on all thinned records against the background. Threshold: 10th-percentile training-presence value (cloglog) used for binary maps.

Null AUC: mean validation AUC of 100 null models built from random points (Bohl et al., 2019); z and p compare the empirical validation AUC with this distribution. Pending: null models still running.

**Table S8.** Long-term (1990–2025) mean predicted suitability by calendar month in Boise, in all surroundings (30 km ring) and in elevation-matched surroundings, with the paired differences (Boise minus surroundings; mean [95% CI] across 36 years).

| Month | Boise | All surroundings | Elevation-matched | Boise − all surroundings | Years > 0 | Adjusted p | Boise − elevation-matched | Years > 0 | Adjusted p |
| --- | --- | --- | --- | --- | --- | --- | --- | --- | --- |
| ***Anax junius*** | | | | | | | | | |
| Jan | 0.164 | 0.140 | 0.160 | 0.0238 [0.0202, 0.0274] | 35 | < 0.001 | 0.0035 [0.0018, 0.0053] | 28 | < 0.001 |
| Feb | 0.248 | 0.206 | 0.244 | 0.0420 [0.0388, 0.0453] | 36 | < 0.001 | 0.0042 [0.0018, 0.0066] | 28 | 0.002 |
| Mar | 0.406 | 0.348 | 0.410 | 0.0584 [0.0535, 0.0633] | 36 | < 0.001 | −0.0038 [−0.0074, −0.0002] | 12 | 0.071 |
| Apr | 0.514* | 0.462 | 0.514 | 0.0522 [0.0472, 0.0572] | 36 | < 0.001 | −0.0003 [−0.0033, 0.0026] | 18 | 0.910 |
| May | 0.584* | 0.564 | 0.588 | 0.0194 [0.0161, 0.0226] | 36 | < 0.001 | −0.0045 [−0.0071, −0.0019] | 11 | 0.003 |
| Jun | 0.652* | 0.634 | 0.653 | 0.0183 [0.0136, 0.0231] | 33 | < 0.001 | −0.0009 [−0.0039, 0.0021] | 17 | 0.661 |
| Jul | 0.639* | 0.648 | 0.645 | −0.0088 [−0.0137, −0.0040] | 10 | < 0.001 | −0.0064 [−0.0092, −0.0035] | 7 | < 0.001 |
| Aug | 0.655* | 0.659 | 0.656 | −0.0048 [−0.0095, 0.0000] | 12 | 0.053 | −0.0012 [−0.0049, 0.0025] | 18 | 0.653 |
| Sep | 0.660* | 0.647 | 0.653 | 0.0133 [0.0089, 0.0178] | 30 | < 0.001 | 0.0066 [0.0032, 0.0100] | 28 | 0.001 |
| Oct | 0.567* | 0.533 | 0.563 | 0.0341 [0.0290, 0.0391] | 36 | < 0.001 | 0.0037 [0.0009, 0.0065] | 25 | 0.021 |
| Nov | 0.316 | 0.274 | 0.310 | 0.0427 [0.0371, 0.0484] | 36 | < 0.001 | 0.0068 [0.0032, 0.0105] | 26 | 0.002 |
| Dec | 0.163 | 0.138 | 0.159 | 0.0249 [0.0215, 0.0283] | 36 | < 0.001 | 0.0042 [0.0023, 0.0061] | 25 | < 0.001 |
| ***Sympetrum corruptum*** | | | | | | | | | |
| Jan | 0.052 | 0.039 | 0.051 | 0.0131 [0.0104, 0.0158] | 35 | < 0.001 | 0.0008 [−0.0004, 0.0020] | 21 | 0.271 |
| Feb | 0.132 | 0.097 | 0.130 | 0.0347 [0.0292, 0.0403] | 36 | < 0.001 | 0.0017 [−0.0004, 0.0037] | 24 | 0.167 |
| Mar | 0.353 | 0.278 | 0.371 | 0.0754 [0.0627, 0.0881] | 36 | < 0.001 | −0.0179 [−0.0250, −0.0108] | 5 | < 0.001 |
| Apr | 0.583* | 0.480 | 0.593 | 0.1031 [0.0950, 0.1112] | 36 | < 0.001 | −0.0102 [−0.0158, −0.0046] | 10 | 0.002 |
| May | 0.656* | 0.628 | 0.674 | 0.0287 [0.0224, 0.0351] | 35 | < 0.001 | −0.0174 [−0.0241, −0.0106] | 6 | < 0.001 |
| Jun | 0.789* | 0.763 | 0.790 | 0.0267 [0.0203, 0.0330] | 34 | < 0.001 | −0.0004 [−0.0046, 0.0037] | 20 | 0.910 |
| Jul | 0.908* | 0.895 | 0.905 | 0.0132 [0.0083, 0.0181] | 32 | < 0.001 | 0.0039 [−0.0002, 0.0080] | 27 | 0.107 |
| Aug | 0.892* | 0.882 | 0.896 | 0.0105 [0.0053, 0.0156] | 29 | < 0.001 | −0.0039 [−0.0082, 0.0004] | 13 | 0.126 |
| Sep | 0.829* | 0.813 | 0.829 | 0.0161 [0.0124, 0.0198] | 33 | < 0.001 | 0.0000 [−0.0037, 0.0037] | 23 | 0.993 |
| Oct | 0.681* | 0.613 | 0.684 | 0.0679 [0.0594, 0.0764] | 36 | < 0.001 | −0.0027 [−0.0085, 0.0031] | 16 | 0.451 |
| Nov | 0.212 | 0.162 | 0.208 | 0.0502 [0.0397, 0.0606] | 36 | < 0.001 | 0.0045 [−0.0013, 0.0103] | 17 | 0.179 |
| Dec | 0.049 | 0.036 | 0.049 | 0.0129 [0.0099, 0.0160] | 36 | < 0.001 | 0.0007 [−0.0005, 0.0018] | 17 | 0.340 |
| ***Tramea lacerata*** | | | | | | | | | |
| Jan | 0.025 | 0.020 | 0.025 | 0.0052 [0.0043, 0.0062] | 35 | < 0.001 | 0.0008 [0.0004, 0.0013] | 29 | 0.001 |
| Feb | 0.037 | 0.029 | 0.036 | 0.0082 [0.0072, 0.0093] | 36 | < 0.001 | 0.0012 [0.0006, 0.0019] | 30 | < 0.001 |
| Mar | 0.068 | 0.054 | 0.067 | 0.0141 [0.0125, 0.0157] | 36 | < 0.001 | 0.0007 [−0.0005, 0.0019] | 21 | 0.340 |
| Apr | 0.116 | 0.093 | 0.115 | 0.0235 [0.0209, 0.0261] | 36 | < 0.001 | 0.0016 [−0.0005, 0.0036] | 24 | 0.190 |
| May | 0.282 | 0.231 | 0.283 | 0.0502 [0.0454, 0.0549] | 36 | < 0.001 | −0.0013 [−0.0048, 0.0022] | 17 | 0.577 |
| Jun | 0.575* | 0.483 | 0.569 | 0.0917 [0.0830, 0.1003] | 36 | < 0.001 | 0.0058 [0.0008, 0.0108] | 23 | 0.045 |
| Jul | 0.728* | 0.706 | 0.729 | 0.0224 [0.0155, 0.0294] | 33 | < 0.001 | −0.0007 [−0.0042, 0.0029] | 17 | 0.816 |
| Aug | 0.715* | 0.700 | 0.713 | 0.0149 [0.0068, 0.0230] | 27 | < 0.001 | 0.0013 [−0.0037, 0.0063] | 19 | 0.712 |
| Sep | 0.533* | 0.465 | 0.505 | 0.0674 [0.0592, 0.0757] | 36 | < 0.001 | 0.0276 [0.0205, 0.0347] | 31 | < 0.001 |
| Oct | 0.166 | 0.138 | 0.157 | 0.0278 [0.0230, 0.0325] | 36 | < 0.001 | 0.0089 [0.0053, 0.0125] | 31 | < 0.001 |
| Nov | 0.053 | 0.044 | 0.051 | 0.0095 [0.0077, 0.0113] | 36 | < 0.001 | 0.0027 [0.0016, 0.0039] | 30 | < 0.001 |
| Dec | 0.026 | 0.020 | 0.024 | 0.0054 [0.0044, 0.0063] | 36 | < 0.001 | 0.0011 [0.0006, 0.0016] | 28 | < 0.001 |
| ***Libellula pulchella*** | | | | | | | | | |
| Jan | 0.061 | 0.058 | 0.060 | 0.0038 [0.0023, 0.0054] | 32 | < 0.001 | 0.0016 [0.0009, 0.0022] | 27 | < 0.001 |
| Feb | 0.089 | 0.077 | 0.085 | 0.0125 [0.0100, 0.0150] | 36 | < 0.001 | 0.0037 [0.0028, 0.0046] | 35 | < 0.001 |
| Mar | 0.191 | 0.160 | 0.190 | 0.0315 [0.0283, 0.0346] | 36 | < 0.001 | 0.0018 [−0.0006, 0.0042] | 21 | 0.186 |
| Apr | 0.332 | 0.283 | 0.332 | 0.0490 [0.0453, 0.0527] | 36 | < 0.001 | 0.0000 [−0.0036, 0.0036] | 13 | 0.993 |
| May | 0.585* | 0.514 | 0.581 | 0.0708 [0.0659, 0.0757] | 36 | < 0.001 | 0.0033 [−0.0001, 0.0066] | 24 | 0.103 |
| Jun | 0.731* | 0.693 | 0.728 | 0.0380 [0.0319, 0.0442] | 36 | < 0.001 | 0.0031 [0.0007, 0.0054] | 29 | 0.024 |
| Jul | 0.720* | 0.717 | 0.714 | 0.0034 [−0.0010, 0.0077] | 20 | 0.136 | 0.0059 [0.0023, 0.0094] | 27 | 0.004 |
| Aug | 0.720* | 0.721 | 0.720 | −0.0015 [−0.0052, 0.0022] | 19 | 0.440 | −0.0001 [−0.0029, 0.0026] | 15 | 0.967 |
| Sep | 0.704* | 0.678 | 0.699 | 0.0256 [0.0215, 0.0297] | 36 | < 0.001 | 0.0045 [0.0022, 0.0068] | 27 | < 0.001 |
| Oct | 0.395 | 0.351 | 0.388 | 0.0437 [0.0379, 0.0495] | 36 | < 0.001 | 0.0070 [0.0032, 0.0108] | 26 | 0.002 |
| Nov | 0.125 | 0.108 | 0.120 | 0.0176 [0.0146, 0.0206] | 36 | < 0.001 | 0.0047 [0.0026, 0.0069] | 30 | < 0.001 |
| Dec | 0.061 | 0.057 | 0.059 | 0.0036 [0.0017, 0.0054] | 29 | < 0.001 | 0.0017 [0.0009, 0.0026] | 26 | < 0.001 |
| ***Pachydiplax longipennis*** | | | | | | | | | |
| Jan | 0.029 | 0.024 | 0.028 | 0.0049 [0.0041, 0.0057] | 35 | < 0.001 | 0.0011 [0.0007, 0.0014] | 29 | < 0.001 |
| Feb | 0.044 | 0.035 | 0.042 | 0.0087 [0.0078, 0.0096] | 36 | < 0.001 | 0.0017 [0.0012, 0.0022] | 33 | < 0.001 |
| Mar | 0.086 | 0.070 | 0.086 | 0.0161 [0.0148, 0.0174] | 36 | < 0.001 | 0.0008 [−0.0002, 0.0018] | 26 | 0.152 |
| Apr | 0.150 | 0.123 | 0.149 | 0.0272 [0.0249, 0.0294] | 36 | < 0.001 | 0.0013 [−0.0003, 0.0029] | 26 | 0.162 |
| May | 0.325 | 0.273 | 0.325 | 0.0524 [0.0481, 0.0567] | 36 | < 0.001 | 0.0005 [−0.0022, 0.0031] | 19 | 0.830 |
| Jun | 0.555* | 0.486 | 0.550 | 0.0695 [0.0626, 0.0763] | 36 | < 0.001 | 0.0050 [0.0018, 0.0081] | 23 | 0.006 |
| Jul | 0.597* | 0.597 | 0.603 | −0.0002 [−0.0078, 0.0074] | 14 | 0.952 | −0.0054 [−0.0077, −0.0031] | 7 | < 0.001 |
| Aug | 0.615* | 0.611 | 0.615 | 0.0036 [−0.0021, 0.0092] | 18 | 0.225 | 0.0002 [−0.0025, 0.0028] | 17 | 0.959 |
| Sep | 0.518 | 0.468 | 0.500 | 0.0501 [0.0449, 0.0554] | 36 | < 0.001 | 0.0183 [0.0148, 0.0219] | 35 | < 0.001 |
| Oct | 0.197 | 0.168 | 0.189 | 0.0290 [0.0254, 0.0326] | 36 | < 0.001 | 0.0074 [0.0047, 0.0101] | 30 | < 0.001 |
| Nov | 0.063 | 0.052 | 0.060 | 0.0101 [0.0085, 0.0117] | 36 | < 0.001 | 0.0028 [0.0017, 0.0039] | 31 | < 0.001 |
| Dec | 0.029 | 0.024 | 0.028 | 0.0051 [0.0043, 0.0060] | 36 | < 0.001 | 0.0013 [0.0009, 0.0018] | 32 | < 0.001 |
| ***Rhionaeschna multicolor*** | | | | | | | | | |
| Jan | 0.048 | 0.034 | 0.049 | 0.0142 [0.0111, 0.0173] | 35 | < 0.001 | −0.0015 [−0.0032, 0.0002] | 10 | 0.131 |
| Feb | 0.096 | 0.068 | 0.097 | 0.0274 [0.0226, 0.0323] | 36 | < 0.001 | −0.0014 [−0.0040, 0.0013] | 16 | 0.389 |
| Mar | 0.186* | 0.145 | 0.203 | 0.0410 [0.0302, 0.0518] | 36 | < 0.001 | −0.0173 [−0.0244, −0.0103] | 6 | < 0.001 |
| Apr | 0.320* | 0.244 | 0.328 | 0.0761 [0.0655, 0.0868] | 36 | < 0.001 | −0.0080 [−0.0164, 0.0005] | 13 | 0.107 |
| May | 0.439* | 0.397 | 0.467 | 0.0416 [0.0339, 0.0493] | 34 | < 0.001 | −0.0281 [−0.0392, −0.0170] | 5 | < 0.001 |
| Jun | 0.706* | 0.642 | 0.706 | 0.0639 [0.0482, 0.0796] | 31 | < 0.001 | −0.0002 [−0.0088, 0.0085] | 21 | 0.993 |
| Jul | 0.810* | 0.813 | 0.809 | −0.0034 [−0.0139, 0.0071] | 17 | 0.530 | 0.0005 [−0.0078, 0.0087] | 21 | 0.965 |
| Aug | 0.815* | 0.815 | 0.827 | −0.0006 [−0.0108, 0.0096] | 12 | 0.917 | −0.0126 [−0.0214, −0.0038] | 9 | 0.012 |
| Sep | 0.758* | 0.711 | 0.745 | 0.0462 [0.0366, 0.0557] | 33 | < 0.001 | 0.0125 [0.0035, 0.0215] | 26 | 0.016 |
| Oct | 0.446* | 0.373 | 0.443 | 0.0735 [0.0624, 0.0846] | 36 | < 0.001 | 0.0032 [−0.0083, 0.0148] | 21 | 0.689 |
| Nov | 0.145 | 0.108 | 0.144 | 0.0370 [0.0281, 0.0459] | 36 | < 0.001 | 0.0006 [−0.0049, 0.0061] | 15 | 0.910 |
| Dec | 0.041 | 0.029 | 0.043 | 0.0126 [0.0097, 0.0154] | 36 | < 0.001 | −0.0021 [−0.0033, −0.0008] | 10 | 0.004 |

* Boise mean at or above the species' 10th-percentile training-presence threshold (Table S7). Years > 0: number of the 36 years in which Boise was more suitable than the comparison area. Adjusted p: one-sample t-test of the 36 yearly differences, with p-values adjusted by the Benjamini–Hochberg procedure across the 72 species-months of each comparison. Suitability values are cover-weighted means of 800 m cells.
