## Supplemental Text for "Sex in the City: Reproductive Behaviors of Dragonflies (Odonata: Anisoptera) across an Urban Landscape"

**Note S1. Species Distribution Modeling**

***Occurrence data from GBIF***

Occurrence data for each species were obtained from the Global Biodiversity Information Facility (GBIF; GBIF.org, 2026a–f). We restricted the search to records from the conterminous United States from 1990–2025 that were recorded as present, had coordinates, had no geospatial issues, and had a coordinate uncertainty of 1 km or less. To reduce sampling bias and spatial autocorrelation, records were thinned within each year and month so that no two records were closer than 20 km, using the R v. 4.5.3 package GeoThinneR (Mestre-Tomás, 2026); year-months with fewer than five records before or after thinning were not used. Exact duplicate records (same species, coordinates, year and month) were removed before thinning, and thinned records in cells without PRISM values were discarded (Table S7).

***Climate predictors***

To generate spatially representative climate predictions for each month, monthly data for 1990–2025 for precipitation (ppt), maximum temperature (tmax), and minimum temperature (tmin) were obtained from the Parameter-elevation Regressions on Independent Slopes Model (PRISM) at a spatial resolution of 800 m (Daly et al., 2008; PRISM Climate Group, 2026). Following the dynamic modeling approach of Tolman et al. (2025), each thinned occurrence record was assigned the climate values of its grid cell in the month and year in which it was observed.

***Background***

When using occurrence data from community-science platforms aggregated by GBIF (e.g., iNaturalist), observation effort often produces biases in the data which result in clustering around accessible paths like roads. Using kernel density estimation in R (ks package; Duong, 2007), we generated a kernel density surface layer from observations of all dragonfly species within our study extent (the conterminous United States; 352,754 records from 1990–2020) to account for sampling bias (Phillips et al., 2009; Fourcade et al., 2014). For each species, we sampled 50,000 background points (with replacement) in proportion to this surface, and assigned each background point a year and month drawn at random in proportion to the number of that species’ thinned occurrence records in each year-month. Each background point was given the PRISM climate values of its location in that month, so that occurrence and background records share the same distribution across years and months. Background points are not treated as absences; they characterize the climatic conditions available across the study extent during the months in which the species was recorded, against which conditions at occurrence records are contrasted during model fitting.

***Model fitting and evaluation***

Species distribution models (SDMs) were fitted separately for each species using the maximum entropy modeling framework implemented by maxnet (Phillips et al., 2006, 2017) through the ENMeval package in R (Kass et al., 2021). Models were based on presence-only occurrence data and background environmental samples, while the three climate predictors were included in the candidate models without additional environmental variables or multicollinearity filtering. We tuned model complexity by testing four feature-class combinations, linear (L), quadratic (Q), linear-quadratic (LQ), and linear-quadratic-hinge (LQH), each with regularization multipliers of 1 through 5 (20 candidate models per species). Candidate models were evaluated with random five-fold cross-validation, using the same folds for all candidate models of a species. The optimal model for each species was the candidate with the highest mean validation AUC (area under the receiver operating characteristic curve; a threshold-independent metric), with ties broken by the lowest mean omission rate of held-out records at the 10th-percentile training-presence threshold (a threshold-dependent metric; Radosavljevic & Anderson, 2014).

For each selected model we report the mean validation AUC; the difference between training and validation AUC (AUCdiff), an index of overfitting; the omission rate of held-out records at the 10th-percentile training-presence threshold (OR10), which should be close to 0.10 for a model that is not overfit (Radosavljevic & Anderson, 2014); and the continuous Boyce index (CBI) of held-out records, calculated with the R package ecospat (Hirzel et al., 2006; Di Cola et al., 2017). We also report the Boyce index of the final model for all thinned records against the background, and checked that the selected setting had the lowest AICc among the candidates (Table S7). To test whether each model performed better than expected by chance, we built 100 null models per species with the ENMnulls function of ENMeval (Bohl et al., 2019). Each null model used the selected feature classes and regularization multiplier, was fitted to the same number of points drawn at random from the pooled occurrence and background records, and was evaluated against the empirical occurrence records withheld in random five-fold cross-validation. The empirical validation AUC was compared with the null distribution with a z-score and one-tailed p-value. Because null points are drawn from a pool that includes the occurrences and are evaluated against the real held-out records, this test is conservative. Response curves were drawn by varying one predictor across its 1st–99th percentile (occurrences and background combined) while holding the other two at their background medians (Fig. S5).

Using the selected model, we generated continuous predictions (complementary log-log [cloglog] output) across Idaho for every month from 1990–2025. For each species, a threshold for the 10th percentile of predicted suitability values was calculated at the thinned occurrence records used to fit the model. Cells with predicted suitability at or above this threshold were classified as suitable (1) and all others as unsuitable (0). We further averaged the monthly predictions across the 36 years within each calendar month to give long-term mean suitability (Figs S6 and S7).

***Projections and the Boise comparison***

We compared monthly climate suitability between the 2020 Census Boise City urban area (US Census Bureau TIGER/Line, accessed with the R package tigris; Walker, 2024) and the surrounding landscape within 30 km of it, excluding all other Census urban areas so that the comparison area was non-urban. Because this ring includes the higher, cooler Boise Foothills, we also compared Boise with only those ring cells whose elevation was within 100 m of Boise’s median elevation (elevation from AWS Terrain Tiles, accessed with the R package elevatr**; Hollister et al., 2023**). We extracted area-weighted mean suitability values from the monthly suitability rasters of the selected models using the R package terra (Hijmans, 2024) and averaged them by calendar month across all years for each species using dplyr (Wickham et al., 2023).

The Boise urban area (366 km²) includes Meridian, Garden City and Eagle. The urban areas removed from the ring were Kuna, Nampa, Emmett, Middleton and Star (224 km²), leaving a 5,338 km² comparison area. Elevation (c. 100 m resolution) was averaged onto the 800 m PRISM grid. The median elevation of Boise cells was 819 m, so the elevation-matched area comprised ring cells between 719 and 919 m (2,252 km², or 42% of the ring; Fig. S4). In all area means, each cell was weighted by the fraction of it lying inside the area.

***Statistical analysis***

For each species, year and month we calculated the difference in mean suitability between Boise and each comparison area (Boise minus surroundings). For each species and calendar month we summarised the 36 yearly differences (1990–2025) by their mean, a 95% confidence interval and the number of years in which Boise was more suitable. We tested whether the mean differed from zero with a one-sample t-test, which is equivalent to a paired t-test of Boise against its surroundings with years as replicates. P-values were adjusted for multiple testing with the Benjamini–Hochberg procedure across the 72 species-months of each comparison (Benjamini & Hochberg, 1995; Table S8). We also averaged the 12 monthly differences within each year and tested the 36 annual values in the same way (Table 4). For each species, the peak month was the calendar month with the highest long-term mean suitability in Boise, and suitable months were those in which long-term mean suitability in Boise was at or above the species' 10th-percentile training-presence threshold. Because consecutive years are not fully independent, these tests describe how consistent the differences are across climate years; they do not include uncertainty in the fitted models.

***Sensitivity analysis***

To check that the conclusions did not depend on how background climate was assigned, we repeated the analysis with the same background locations but with each point given its climate averaged over all 432 months of 1990–2025, instead of the climate of a single year-month. We compared Boise's seasonal suitability and its difference from elevation-matched surroundings between the two runs (Fig. S8).
